# SpatialRNA: a python package for easy application of Graph Neural Network models on single-molecule spatial transcriptomics dataset

**DOI:** 10.1101/2025.03.20.644258

**Authors:** Ruqian Lyu, Annika Vannan, Jonathan A. Kropski, Nicholas E. Banovich, Davis J. McCarthy

## Abstract

Image-based spatial transcriptomics deliver gene expression measurements of RNA transcripts in tissue slices with single-molecule resolution and spatial context preserved. Modern Graph Neural Network (GNN) models are promising methods for capturing the complex molecular and cellular phenotypes in tissues at single-transcript and single-cell levels. A key application of GNNs is the detection of spatial domains or niches, that is, groups of molecules and/or cells that collaboratively work together to produce complex phenotypes. Due to the vast number of detected transcripts in image-based spatial transcriptomics, applying GNNs on RNA molecule graphs is not trivial. We present a Python package, SpatialRNA, for easy (sub)graph generation from tissue samples and provide comprehensive tutorials for convenient and efficient application of Graph Neural Network models under the PyG framework. Availability and implementation: The SpatialRNA package is freely accessible from GitLab https://gitlab.svi.edu.au/biocellgen-public/spatialrna and can be installed via pip. Python notebooks and scripts used in case studies are freely accessible at https://gitlab.svi.edu.au/biocellgen-public/case_study_ipf and https://gitlab.svi.edu.au/biocellgen-public/case_study_xenium_5k_panel.

## Introduction

Image-based spatial transcriptomics (iST) platforms are able to detect RNA transcripts within the spatial context of tissues, allowing quantification of gene expression in cells and tissues in a spatially-aware manner. These platforms, such as MERFISH (Vizgen) (1), seqFISH (2, 3), and Xenium (10x Genomics), commonly build upon Fluorescence in situ hybridization (FISH) techniques that visualise RNA locations through florescent-dye-attached probes designed to specifically bind to transcript targets. With iST platforms advancing, hundreds to thousands of gene species are measured in parallel in a given tissue sample, yielding high-throughput single-molecule resolution spatial gene expression data.

Broadly, there are two analysis approaches for iST data: cell-based and cell-free approaches (4, 5). With cell-based approaches, cell segmentation (6–8) is required to obtain cell boundaries using auxiliary imaging data such as nuclei and cell membrane staining as well as gene expression profiles. Cell-level gene expression is then derived based on cell boundaries through summarising molecules within cell boundaries, which allows application of many existing single-cell analysis methods. Performance of cell-based analysis is heavily influenced by the quality of cell segmentation, and errors in cell segmentation introduce biases in the final outcome. Cell-free analysis methods perform modelling directly on the detected transcripts and their spatial neighbourhoods for cell or tissue typing (4, 5). Cell-free methods are thus more suitable for iST data without reliable cell segmentations, for example, iST data without quality membrane staining images, or tissues with cells that are particularly hard to segment due to irregular shapes or high cell density.

Graph neural network (GNN) models are deep machine learning methods that are designed to work with graph structured data. These methods also show great potential in cellfree iST data analysis, with specific GNN models such as GraphSAGE (9) having been successfully applied to spatial RNA graphs for learning diverse spatial organisation of RNA transcript neighbourhoods (i.e., spatial domains or niches) (10, 11). Characterising spatial domains or niches in the tissue is helpful for understanding mechanisms of how groups of cells work together to perform complex functions and essential for associating disease phenotypes with coordinated changes in molecular and cellular compositions. To apply GNNs on iST data in a cell-free (molecule-based) manner, spatial RNA graphs are constructed using RNA molecules as nodes and edges are added to connect RNA molecules to their spatial neighbours. Input nodes in the graph are associated with an initial feature vector, usually a one-hot-encoding representation of the gene labels. GNNs are then deployed to learn functions of aggregating neighbourhoods of individual molecules to project each molecule onto a latent space where molecules with similar neighbourhood compositions are closely embedded in similar locations in the latent space. However, applying GNNs on RNA molecule graphs, especially in large tissue samples (both large in physical dimension of each tissue sample and/or large in sample size), is not trivial due to the vast number of RNA molecules detected, at the scale of millions of transcripts. We recently introduced our working GNN computational workflow in a study that investigated pulmonary fibrosis using a total set of 45 lung tissue samples (11), together containing about 299 million RNA transcripts. The workflow consisted of multiple steps with designated Python scripts for graph construction, sub- graph sampling, and model training, and it was able to detect spatial tissue domains across healthy and fibrotic lungs in a cell-free manner. To make the application of GNNs on such iST data easier and more streamlined, we hereby introduce a python package `SpatialRNA`, that modularises each processing component and enables convenient application of a range of existing or novel GNN architectures on iST samples utilising the modern GNN library PyG (https://github.com/pyg-team/pytorch_geometric).

## Methods and implementations

The goal of applying GNNs for cell-free molecule-based analysis for iST data for spatial domain identification is to learn new representations of individual molecules. Specifically, the GNN assigns a latent embedding vector for each individual molecule based on the composition of its spatial neighbourhood (defined by a distance threshold). The latent embeddings of molecules can be subsequently used for unsupervised clustering to detect spatial tissue domains across samples (Fig. 1). Since the aim is to identify spatial tissue domains in an unsupervised fashion, there are no classification labels available for the nodes in the graph. Therefore, GNNs can not be trained in a node classification setup. Instead, the GNNs models are trained by solving a link (or edge) classification task. This task involves predicting whether a link exists between a pair of nodes based on the similarity of the profiles of their spatial neighbours, which is supervised by the graph structure (9–11).

**Fig. 1.**
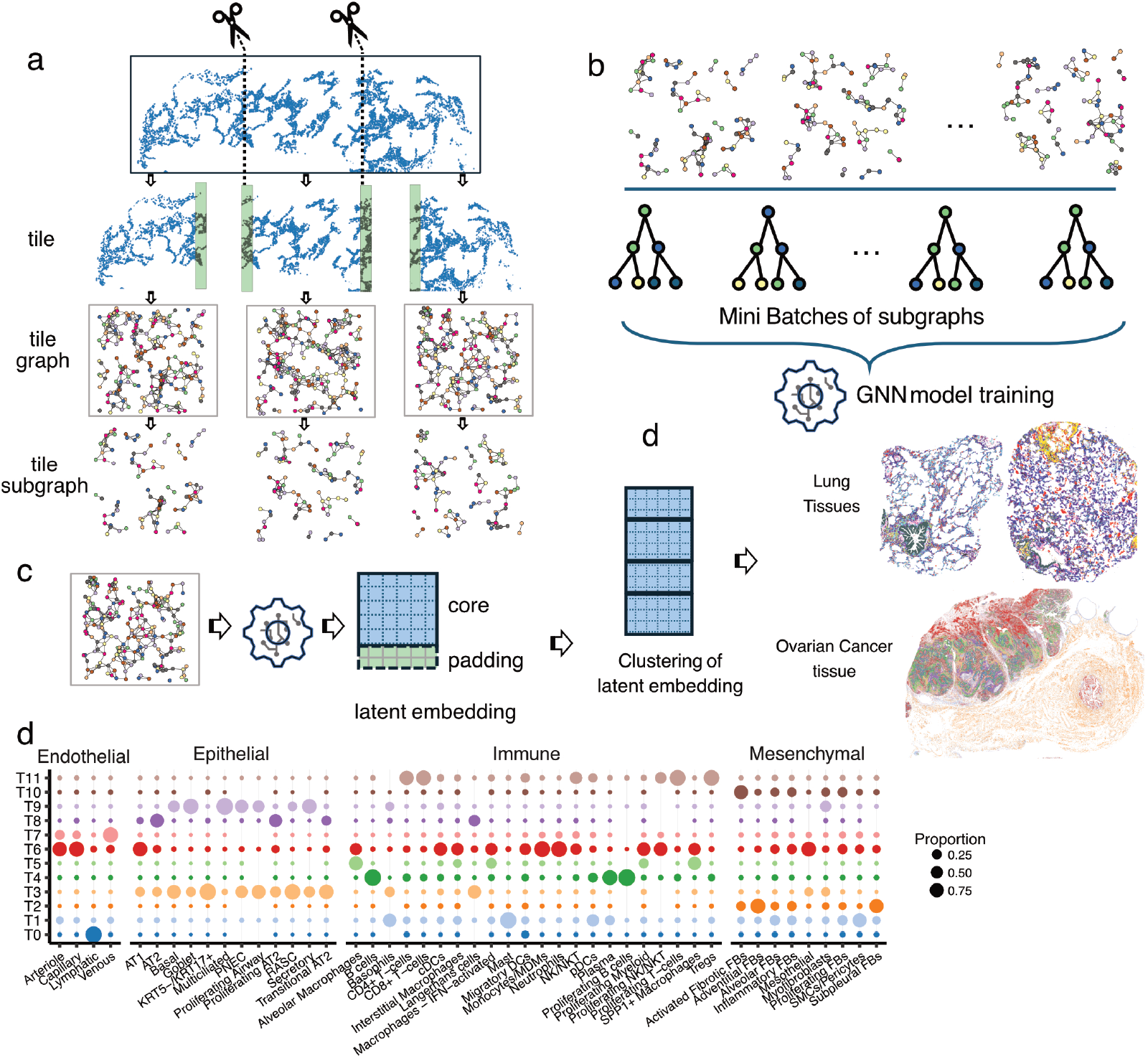
SpatialRNA for easily tiling large tissue samples, constructing tile (sub)graphs, and facilitating mini-batched GNN model training. a) An example tissue is tiled into three tissue tiles. Each tile consists of a core tissue area as well as padding areas to include neighbours of transcripts located on the edges of the tiles. Subgraphs are sampled from the tile graph, and all subgraphs from multiple tissues (tissue tiles) were merged for model training as shown in b. b) Mini-batches of graphs were generated from the merged graph for GNN (e.g., GraphSAGE, GAT, GATv2) model training. c) Nodes (i.e., transcripts) in each tissue tile graph were projected onto the latent space using the trained GNN model. Nodes in the padding areas are discarded. d) Nodes from multiple tiles of tissues were stacked and clustering was applied to the embedding matrix for partitioning the transcripts into groups. Two lung tissues from case study 1, and the ovarian cancer tissue from case study 2 were visualised after hex bin aggregation. d) The cellular composition of the 12 identified molecular niches (T0–T11) from the idiopathic pulmonary fibrosis (lung) case study 1.

The input data files are the list of individual transcripts with spatial locations in each tissue sample. Starting with the list of spatially detected transcripts, the following major steps are involved in the process: 1) constructing spatial RNA graphs based on a selected radius threshold for each sample; 2) sampling subgraphs from the RNA graphs per sample; 3) merging subgraphs across samples and training of the selected GNN model using the merged graph; and 4) inference of latent embedding for transcripts in individual samples. In the unified embedding space, transcript-based clusters (or niches) can be detected through a selected clustering algorithm. Utilising the PyG framework, we implemented the `SpatialRNA` package that provides convenient construction and processing of graphs, facilitating large-scale iST data analysis.

### Build sample graphs

Spatial RNA graphs are constructed using a radius-based approach for each sample using individual RNA transcripts as nodes. Edges are added between nodes with distances smaller than the selected radius threshold. For large tissue samples when the entire tissue graph does not fit in memory, tiles can be created along the chosen axis, such as along the *x* or *y* axis. The number of tiles is set by the user and each tile is also padded with extra margins to include the required spatial neighbours of transcripts located on the edge of the tile boundaries (to avoid artificial edge effects). A binary label is added to record which transcripts are from the core area or the padded area. Initialisation of the input feature vector for each node/transcript is set as the integer gene ID, which helps to reduce storage costs of the constructed graph. In the training and inference stages, integer IDs are mapped to one-hot-encoding vectors per batch of data.

### Build sample subgraphs

Subgraphs are sampled from each sample graph using the ‘LinkNeighborLoader’ from the PyG library in either a node-based or link-based manner. So-called “seed node-based mode” samples a fixed number of seed nodes first and a fixed number of random walks are created from seed nodes to reach the first layer neighbours. This mode was first adopted in the Node2Vec model (12). Nodes reached by these walks are included in the subgraph along with their neighbours that are within a certain number of “hops”. Here, “hop” refers to the distance to a neighbour measured by the number of steps required to reach the neighbour. Alternatively, “link-based subgraph sampling” randomly samples a chosen number of edges, and nodes reached by these edges along with their neighbours within a certain number of hops are included.

As processing each tile or sample is independent from another, tile- or sample-based processes can be run in parallel (i.e., in multiple separate processing jobs), which includes the sample graph and subgraph construction, and embedding inference.

### Model training

PyG library offers predefined ready-to-use GNN layers and GNN models (with in-built GPU acceleration supported through the PyTorch API) such as Graph-SAGE (9), Graph Attention Networks (GAT) (13), and GATv2 (14), making it convenient to experiment with different architectures as well as construct customised GNNs. Once the GNN model is configured, the model is trained using the merged subgraphs across tiles and samples. Minibatching from the joined graph is performed for model training via edge label prediction.

### Inference on embeddings of individual transcripts

Inference on embeddings is performed for all nodes in the tile graph (in a mini-batch manner) but only nodes in the core area of each tile are included in the final output embedding matrix. The embedding matrices from multiple tiles originating from one tissue sample can be stacked to derive the embedding matrix for the full list of transcripts in the sample.

## Case studies

To demonstrate the usage of `SpatialRNA` we conducted two case studies analysing spatially detected molecules from both lung tissues and a ovarian caner tissue generated from the Xenium platform (10x Genomics).

### Case study 1: Human pulmonary fibrosis dataset

We obtained the list of detected transcripts from 343 genes in 45 lung tissues processed with 10x Xenium from GEO with accession number GSE250346 (11). We then processed each sample individually (without tiling) and create the sample graph using a radius threshold of 3.0 (microns). We subsequently sampled subgraphs from individual samples in a node-based fashion, which sampled 5,000 seed nodes and performed 5 random walks per seed node for positive edge sampling. Nodes in the set of sampled edges along with 2-hop neighbours were kept in the subgraph. Subgraphs were then joined for GNN training. We constructed a 2-layer GAT (13) model as implemented in torch.geometrics (v2.6.1) with L2 normalisation at each layer, and trained the model using the joined subgraphs. Using the trained GNN model, we predicted the embeddings of nodes (individual transcripts) across 45 tissue samples. Lastly, we jointly clustered transcripts across all samples using their embeddings with a Gaussian Mixture model. Jupyter notebooks are available in the Supplementary Material as well as in a GitLab repository. To demonstrate that spatial niches capture multicellular structure, we obtained the cell type labels (11) and calculated the cell type proportion assigned into each identified molecular niche (Fig. 1.d).

### Case study 2: Human ovarian cancer sample with 5K gene panel

As a demonstration on state-of-art large-scale iST data, we analysed the human ovarian cancer sample processed on Xenium with a panel of 5,000 genes from 10x Genomics using our method. Using `SpatialRNA` we first tiled the tissue to 30 tiles along the *y* axis. We created RNA spatial graphs and subgraphs for individual tiles. Subgraphs were generated in a link-based manner where 5,000 links were sampled first, and nodes in the set of sampled edges along with 2-hop neighbours were kept in the subgraph. We joined the subgraphs from every other tiles (15 tiles were joined) for model training. We then obtained molecule-based niches the same way as for the lung samples and compared the molecule niches with the cell group labels obtained from the reference source (Figs. S3-S6) revealing that the molecule niches recover regions of tumour and non-tumour tissues. Python notebooks are available in the Supplementary Material as well as online on GitLab.

## Conclusion

We presented the open-source python package `SpatialRNA` for convenient and efficient processing of spatially detected transcripts from iST data, which allows seamless integration with GNN methods in the PyG framework, making molecule-based analysis ready for ongoing GNN advancement. We demonstrate the usage of the package and the analysis workflow through two case studies that analysed large-scale iST datasets both in terms of number of samples as well as large tissue area with expanded gene panels. We showed that the molecule-based niches are biologically meaningful through correlating with cell-based results, which also provides biological interpretation for the spatial niches. When there is no cell-based analysis, the overrepresented genes in each niche can be identified and used for interpreting the identified niches along with spatial organisation. While both case studies were conducted on datasets from the Xenium platform, the method generalises to other datasets with a similar format, that is features (e.g., mRNA or protein) with spatial locations. Additionally, the radius-based graph construction can be readily extended to 3D space. The package is currently designed to focus on molecule-based inputs, but can be applied to cell-based inputs with adaptation. Lastly, it is expected to tune parameter configurations for specific datasets, aligning with each specific analysis goals. For examples, the radius thresholds used for constructing graphs control the size of spatial neighbourhoods for spatial smoothing, and increasing radius increases the level of spatial smoothing. The size of the GNN model (e.g., hidden layer sizes) and the size of sample neighbours are also expected to increase with increasing complexity in the dataset.

## Supporting information

Supplementary_case_study1

Supplementary_case_study2

## Author contributions statement

R.L implemented the package, conducted the analysis, and wrote the manuscript. A.V., J.A.K, and N.E.B. generated data. A.V., J.A.K, N.E.B, and D.J.M guided and supervised the analysis. A.V., J.A.K., N.E.B, and D.J.M reviewed the manuscript.

## Acknowledgments

This work was supported by funding from the US National Institutes of Health, R01HG011886 (NEB/DJM) and R01HL145372 (NEB/JAK), and the Australian National Health and Medical Research Council, GNT1195595 (DJM).

## Supplementary Material

The compiled notebooks for two cases studies are available in Supplementary Material case study IPF and case study ovarian cancer.

## References

1. Kok Hao Chen, Alistair N Boettiger, Jeffrey R Moffitt, Siyuan Wang, and Xiaowei Zhuang. RNA imaging. spatially resolved, highly multiplexed RNA profiling in single cells. Science, 348(6233):aaa6090, April 2015. ISSN 0036-8075,1095-9203. doi: 10.1126/science.aaa6090.

2. Eric Lubeck, Ahmet F Coskun, Timur Zhiyentayev, Mubhij Ahmad, and Long Cai. Single-cell in situ RNA profiling by sequential hybridization. Nature methods, 11(4):360–361, April 2014. ISSN 1548-7091,1548-7105. doi: 10.1038/nmeth.2892.

3. Chee-Huat Linus Eng, Michael Lawson, Qian Zhu, Ruben Dries, Noushin Koulena, Yodai Takei, Jina Yun, Christopher Cronin, Christoph Karp, Guo-Cheng Yuan, and Long Cai. Transcriptome-scale super-resolved imaging in tissues by RNA seqFISH. Nature, 568 (7751):235–239, April 2019. ISSN 0028-0836,1476-4687. doi: 10.1038/s41586-019-1049-y.

4. Yichen Si, Changhee Lee, Yongha Hwang, Jeong H Yun, Weiqiu Cheng, Chun-Seok Cho, Miguel Quiros, Asma Nusrat, Weizhou Zhang, Goo Jun, Sebastian Zöllner, Jun Hee Lee, and Hyun Min Kang. FICTURE: scalable segmentation-free analysis of submicron-resolution spatial transcriptomics. Nature methods, 21(10):1843–1854, October 2024. ISSN 1548-7091,1548-7105. doi: 10.1038/s41592-024-02415-2.

5. Yichun He, Xin Tang, Jiahao Huang, Jingyi Ren, Haowen Zhou, Kevin Chen, Albert Liu, Hailing Shi, Zuwan Lin, Qiang Li, Abhishek Aditham, Johain Ounadjela, Emanuelle I Grody, Jian Shu, Jia Liu, and Xiao Wang. ClusterMap for multi-scale clustering analysis of spatial gene expression. Nature communications, 12(1):5909, October 2021. ISSN 2041-1723. doi: 10.1038/s41467-021-26044-x.

6. Carsen Stringer, Tim Wang, Michalis Michaelos, and Marius Pachitariu. Cellpose: a generalist algorithm for cellular segmentation. Nature methods, 18(1):100–106, January 2021. ISSN 1548-7091,1548-7105. doi: 10.1038/s41592-020-01018-x.

7. Yuxing Wang, Wenguan Wang, Dongfang Liu, Wenpin Hou, Tianfei Zhou, and Zhicheng Ji. GeneSegNet: a deep learning framework for cell segmentation by integrating gene expression and imaging. Genome biology, 24(1):235, October 2023. ISSN 1465-6906. doi: 10.1186/s13059-023-03054-0.

8. Xiaohang Fu, Yingxin Lin, David M Lin, Daniel Mechtersheimer, Chuhan Wang, Farhan Ameen, Shila Ghazanfar, Ellis Patrick, Jinman Kim, and Jean Y H Yang. BIDCell: Biologically-informed self-supervised learning for segmentation of subcellular spatial transcriptomics data. Nature communications, 15(1):509, January 2024. ISSN 2041-1723,2041-1723. doi: 10.1038/s41467-023-44560-w.

9. William L Hamilton, Rex Ying, and Jure Leskovec. Inductive representation learning on 4 | bioRχiv Lyu et al. | SpatialRNA large graphs. arXiv [cs.SI], June 2017.

10. Gabriele Partel and Carolina Wählby. Spage2vec: Unsupervised representation of localized spatial gene expression signatures. The FEBS journal, 288(6):1859–1870, March 2021. ISSN 1742-464X,1742-4658. doi: 10.1111/febs.15572.

11. Annika Vannan, Ruqian Lyu, Arianna L Williams, Nicholas M Negretti, Evan D Mee, Joseph Hirsh, Samuel Hirsh, Niran Hadad, David S Nichols, Carla L Calvi, Chase J Taylor, Vasiliy V Polosukhin, Ana P M Serezani, A Scott McCall, Jason J Gokey, Heejung Shim, Lorraine B Ware, Matthew J Bacchetta, Ciara M Shaver, Timothy S Blackwell, Rajat Walia, Jennifer M S Sucre, Jonathan A Kropski, Davis J McCarthy, and Nicholas E Banovich. Spatial transcriptomics identifies molecular niche dysregulation associated with distal lung remodeling in pulmonary fibrosis. Nature genetics, pages 1–12, February 2025. ISSN 1061-4036,1546-1718. doi: 10.1038/s41588-025-02080-x.

12. Aditya Grover and Jure Leskovec. Node2vec: Scalable feature learning for networks. arXiv [cs.SI], July 2016.

13. Petar Veličkovicć, Guillem Cucurull, Arantxa Casanova, Adriana Romero, Pietro Liò, and Yoshua Bengio. Graph attention networks. arXiv [stat.ML], October 2017.

14. Shaked Brody, Uri Alon, and Eran Yahav. How attentive are graph attention networks? arXiv [cs.LG], May 2021.

