## Supplemental_figures for "SpatialRNA: a python package for easy application of Graph Neural Network models on single-molecule spatial transcriptomics dataset"

### Supplementary figures

March 19, 2025

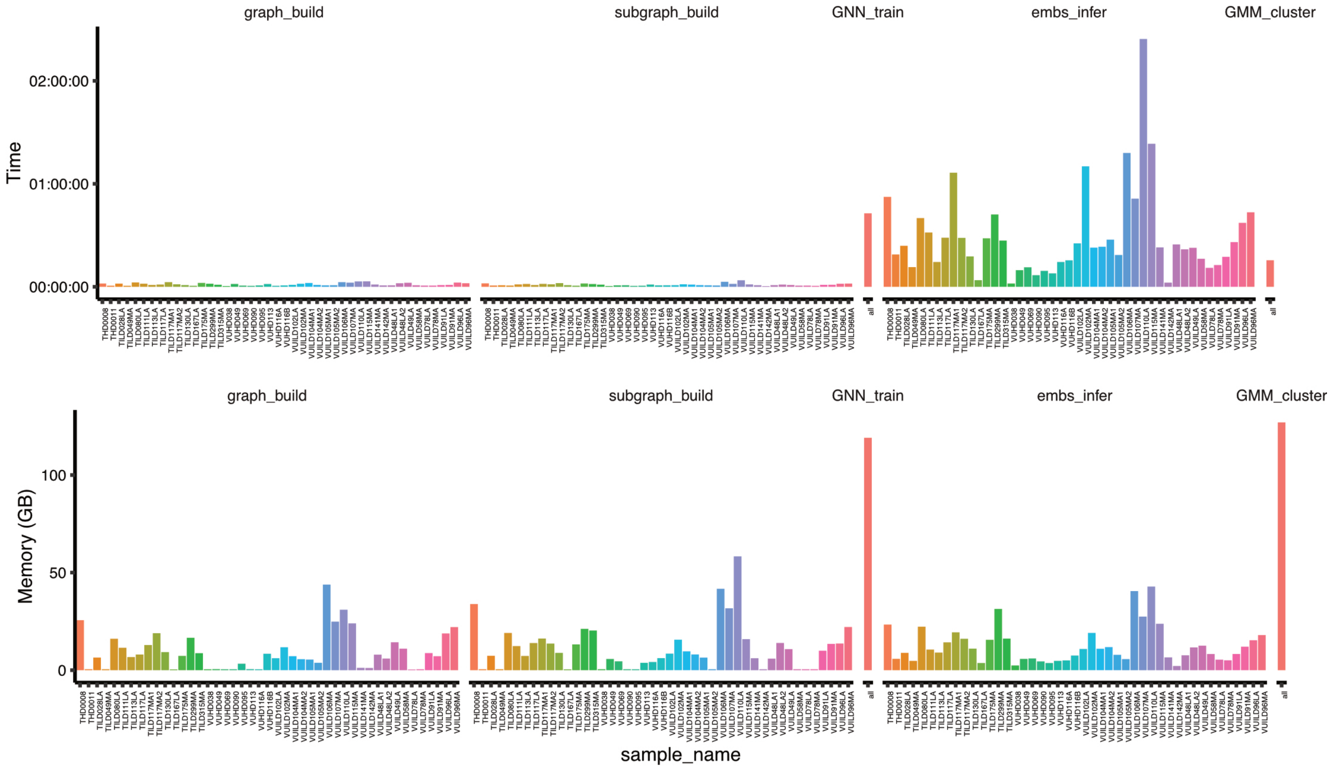


Fig. S1: Running time and RAM usage for each computational process for each sample. The joined subgraphs of individual samples was used for GNN training.

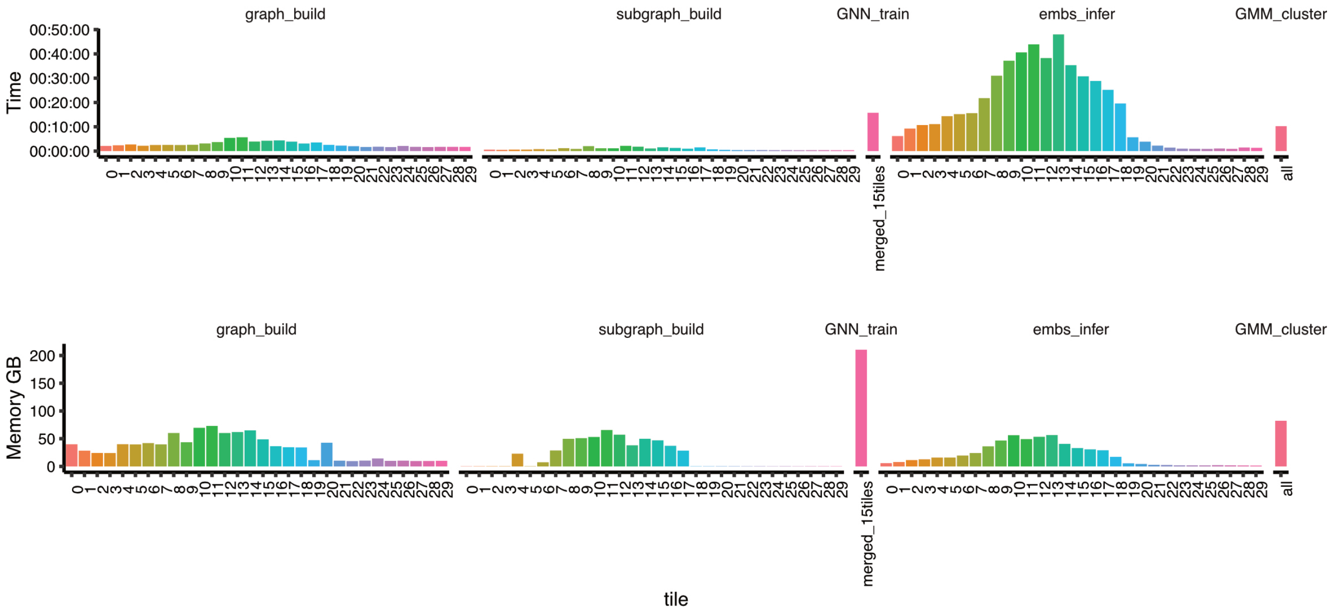


Fig. S2: Running time and RAM usage for each computational process for each sample or joined samples.

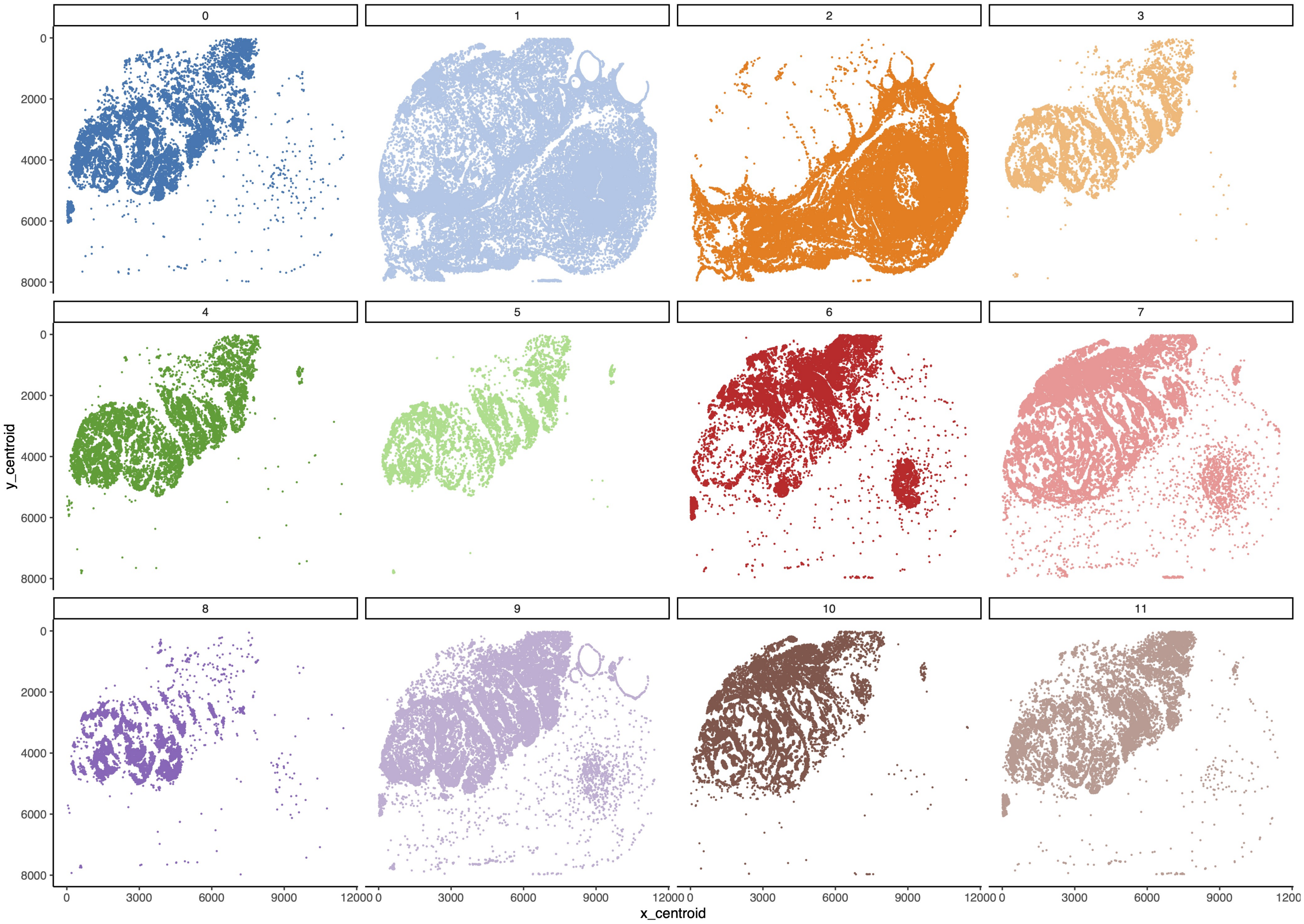


Fig. S3: The twelve identified molecule niches in a ovarian cancer tissue in case study 2. The molecule-based niches were identified as demonstrated in case study 2 and the molecules were hex bin aggregated using bin width 5 (microns). Each hex bin is labelled with the major cluster labels among the spatially residing molecules in each hex bin. To compare with the cell annotations downloaded from 10x Genomics website, we obtained the cell centroids and assigned cell centroids to the hex bins by assigning cell centroids to their closest hex bins. We then transferred the hex bin label to cells.

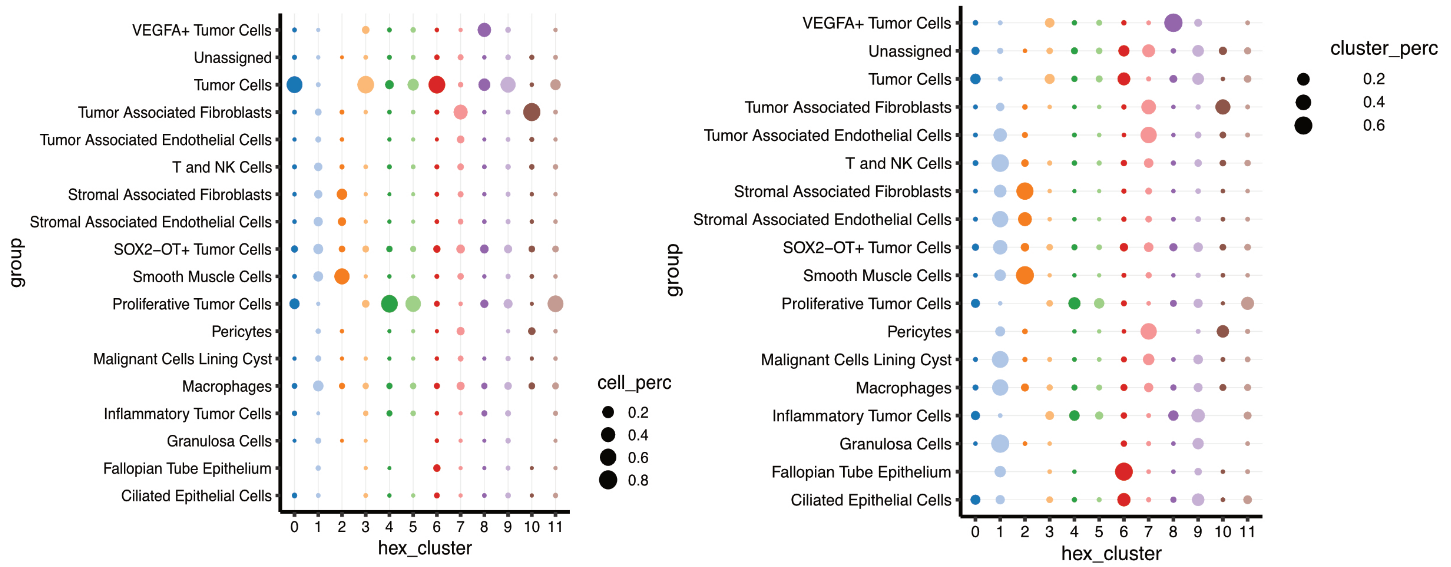


Fig. S4: Relations of cell annotation groups with molecule-based niches. Left panel shows the cell group composition for each molecule niche. Right panel shows the niche composition for each cell group. 'Tumor Cells' were mostly divided into niches 0, 3, and 6.

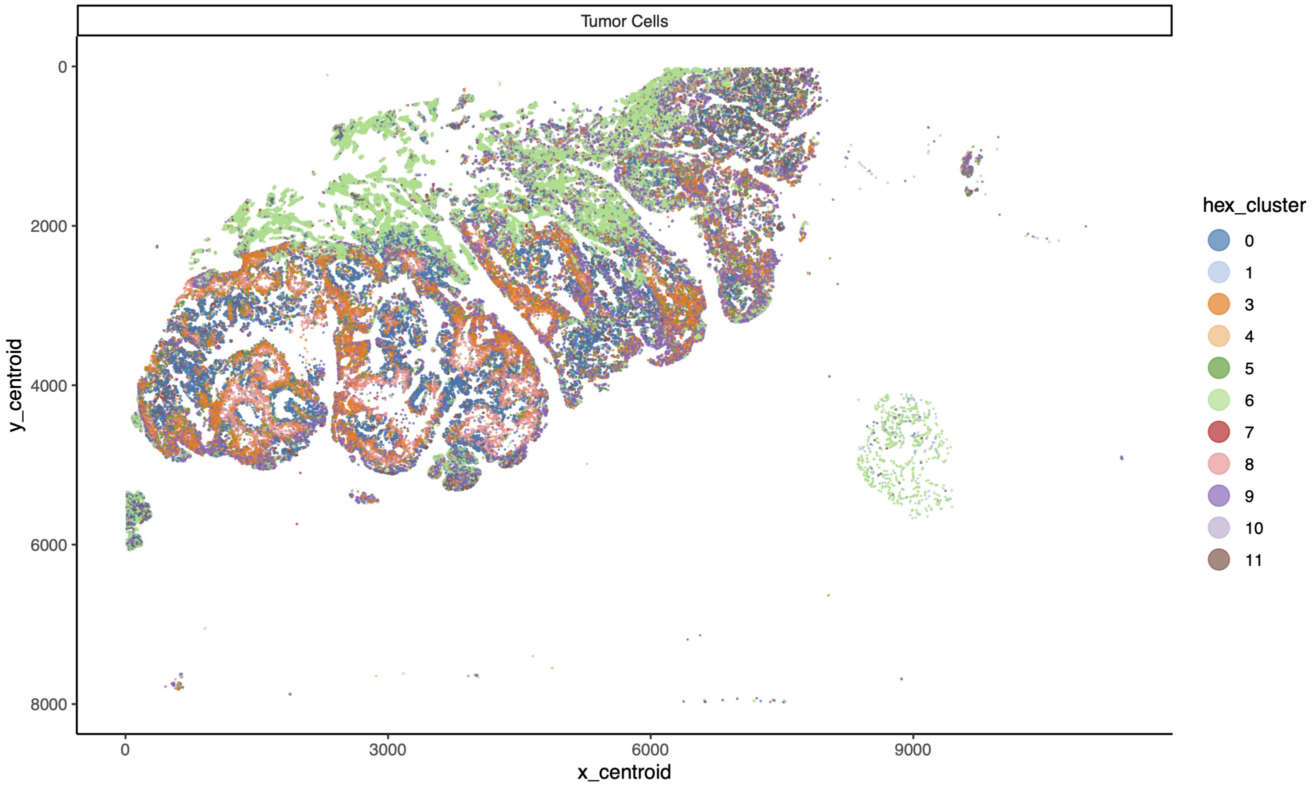


Fig. S5: Visualising 'Tumor Cells' only and colored by their assigned molecule based niche labels.

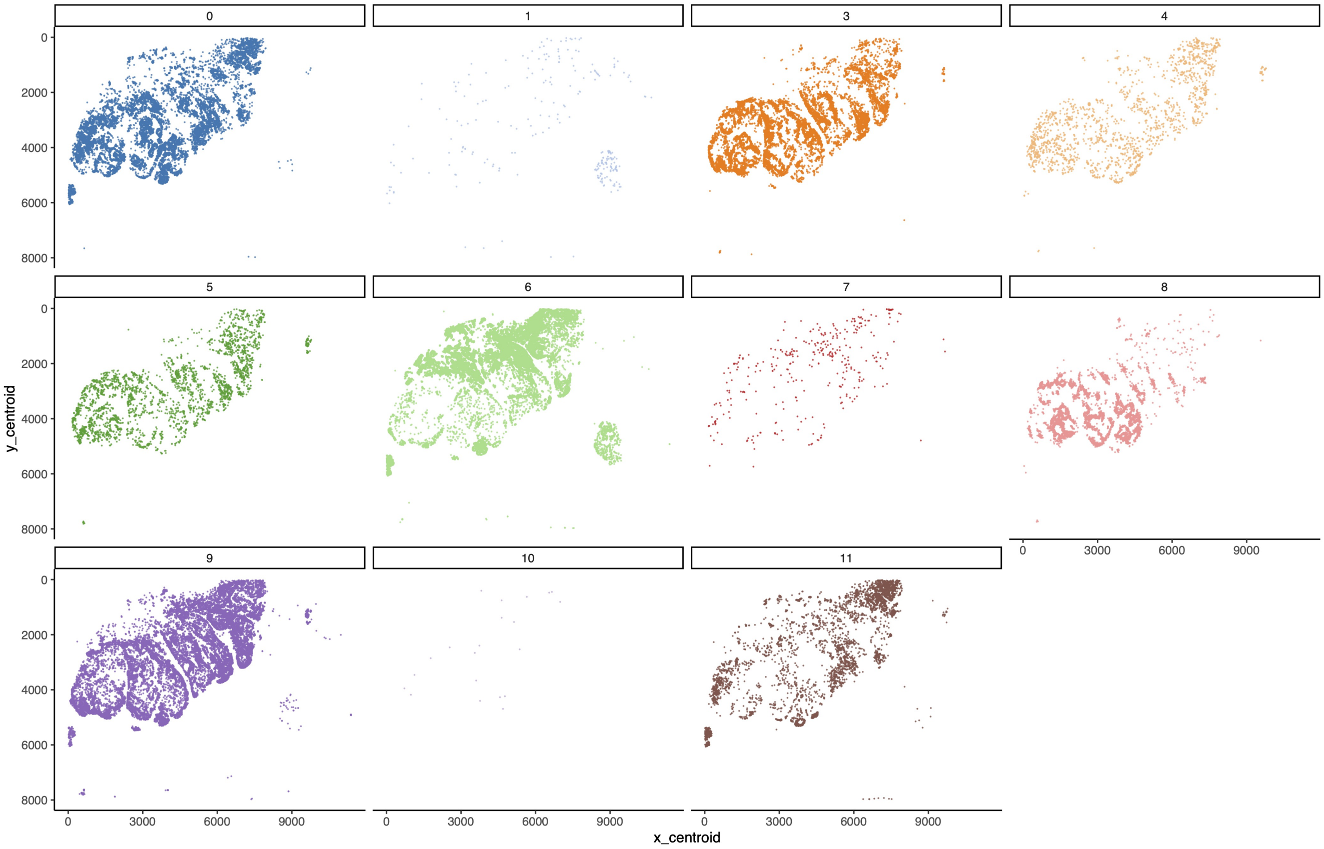


Fig. S6: Visualising 'Tumor Cells' only and coloured by their assigned molecule-based niche labels.
