## Supplementary_case_study1 for "SpatialRNA: a python package for easy application of Graph Neural Network models on single-molecule spatial transcriptomics dataset"

### case_study_ipf_lungs

March 19, 2025

**3 Plot spatial tissue domains in lung samples** **12**

### 1 Lung tissue samples

The Xenium assayed lung tissue samples were obtained from GSE250346 that consists of 45 samples with healthy and fibrosis phenotype. A total of 343 genes were measure for these samples that are of size 3-5 mm in diameter, detecting about 299 million transcripts.

We will analyse transcripts from all 45 samples integrately, through training a GNN model using the joined subgraphs from all samples. We build graphs and subgraphs for samples individually. The following code chunks demonstrate the code applied for one sample. The workflow apply the same processing for 45 samples.

[1]:
```
ls ../workflows/run_generate_subg.smk
```

```
../workflows/run_generate_subg.smk
```

[1]:
```
sample_name = "VUHD038"
```

[2]:
```
import spatialrna
SpatialRNA = spatialrna.SpatialRNA
```

[3]:
```
?SpatialRNA
```

```
Init signature:
SpatialRNA(
    root: str,
    sample_name: Optional[str] = None,
```

```
    radius_r: float = 3.0,
    dim_x: str = 'X',
    dim_y: str = 'Y',
    tile_by_dim: str = 'Y',
    num_tiles: int = 1,
    process_mode: str = 'tile',
    load_type: str = 'tile',
    load_tile_id: int = 0,
    pad_hops: float = 2,
    feature_col='gene',
    max_num_neighbors: int = 500,
    process_tile_ids: List[int] = None,
    one_hot_encoding: dict = None,
    subgraph_mode: str = 'link_based',
    num_sampled_edges: int = 5000,
    num_seed_nodes: int = 5000,
    num_walks: int = 5,
    transform=None,
    pre_transform=None,
    force_reload: bool = False,
    force_resample: bool = False,
    log: bool = True,
    seed: int = 100,
    **kwargs,
) -> None
Docstring:
A spatial RNA graph dataset. One dataset per tissue sample, and optionally creates
tiles from a tissue area. For each generated data.pt, it contains the nodes in␣
 ↪core areas for
making inference for transcripts with complete neighbourhoods.

Args:
    root (str): Root directory where the dataset should be saved.
    sample_name (str): The name of this sample. Appended to the processed␣
 ↪[sample_name]_data.pt.
                       Using basename of root dir if not supplied.
    radius_r (float): The radius for building RNA graphs.
    tile_by_dim (str): Along which dimension to make tiles. It should be a column␣
 ↪in the transcript data.frame.
    num_tiles (int): The number of tiles to create from this tissue sample.
    process_mode (str): The mode of process can be either "tile" or "subgraph".
    load_type (str): If :obj:`"tile"`, load the tile graph data.
        If :obj:`"subgraph"`, load the tile subgraph data.
    load_tile_id (int): Select one tile to load.
    process_tile_ids (List[int]): Select tiles to process including generating␣
 ↪tile graph and subgraph.
```

```
    one_hot_encoding (dict): Dictionary that mapps gene names to integer (or␣
↪one-hot-encoding tensors).
        It is essential to apply the same dictionary for all samples that are␣
↪analyzed together.
    transform (callable, optional): A function/transform that takes in an
        :obj:`torch_geometric.data.Data` object and returns a transformed
        version. The data object will be transformed before every access.
        (default: :obj:`None`)
    pre_transform (callable, optional): A function/transform that takes in
        an :obj:`torch_geometric.data.Data` object and returns a
        transformed version. The data object will be transformed before
        being saved to disk. (default: :obj:`None`)
    force_reload (bool, optional): Whether to re-process the dataset.
        (default: :obj:`False`)
    force_resample (bool, optional): Whether to re-process the tile dataset to␣
↪generate subgraphs.
        (default: :obj:`False`)
    subgraph_mode (str): The mode of subgraph generation can be either␣
↪"link_based" or "node_based".
        (default: :obj:`link_based`)
    num_sampled_edges (int): sample this many edge_index as positive edges when␣
↪generating subgraph from a tile (used in LinkNeighborLoader).
    num_seed_nodes:(int, optional): Number of seed nodes for generating subgraphs.
        (default: :obj:`5000`)
    num_walks:(int, optional): Number of walks for generating subgraphs.
        (default: :obj:`5`)

    log (bool, optional): Whether to print any console output while
        downloading and processing the dataset. (default: :obj:`True`)
    **kwargs (optional): Additional arguments of
        :class:`torch.utils.data.LinkNeighborLoader`, such as :obj:`num_neighbors`
File:           /mnt/beegfs/mccarthy/general/backed_up/rlyu/Projects/spatialrna/
↪spatialrna/spatialrna.py
Type:           type
Subclasses:
```

[2]:

```
import pandas as pd
import numpy as np
import torch
import torch.nn.functional as F
gene_panel = pd.read_csv("../resources/xenium_gene_panel.csv")
gene_panel
```

[2]:

| | Unnamed: 0 | x |
|---|---|---|
| 0 | 1 | EPAS1 |
| 1 | 2 | EMG1 |
| 2 | 3 | MYC |

```
3              4  S100A9
4              5   LPAR1
..           ...     ...
338          339    TERT
339          340    CD27
340          341   KRT17
341          342   PDCD1
342          343  VPREB3

[343 rows x 2 columns]
```

[5]:
```
gene_list = np.unique(gene_panel.x)

x = torch.tensor(np.arange(gene_list.shape[0]))

one_hot_encoding = dict(zip(gene_list, F.one_hot(x, num_classes=gene_list.
 ↪shape[0])))

for k in one_hot_encoding.keys():
    one_hot_encoding[k] = one_hot_encoding[k].double()
print(one_hot_encoding["CD3D"])
```

```
tensor([0., 0., 0., 0., 0., 0., 0., 0., 0., 0., 0., 0., 0., 0., 0., 0., 0., 0.,
        0., 0., 0., 0., 0., 0., 0., 0., 0., 0., 0., 0., 0., 0., 0., 0., 0., 0.,
        0., 0., 0., 0., 0., 0., 0., 0., 0., 0., 0., 0., 0., 0., 0., 1., 0., 0.,
        0., 0., 0., 0., 0., 0., 0., 0., 0., 0., 0., 0., 0., 0., 0., 0., 0., 0.,
        0., 0., 0., 0., 0., 0., 0., 0., 0., 0., 0., 0., 0., 0., 0., 0., 0., 0.,
        0., 0., 0., 0., 0., 0., 0., 0., 0., 0., 0., 0., 0., 0., 0., 0., 0., 0.,
        0., 0., 0., 0., 0., 0., 0., 0., 0., 0., 0., 0., 0., 0., 0., 0., 0., 0.,
        0., 0., 0., 0., 0., 0., 0., 0., 0., 0., 0., 0., 0., 0., 0., 0., 0., 0.,
        0., 0., 0., 0., 0., 0., 0., 0., 0., 0., 0., 0., 0., 0., 0., 0., 0., 0.,
        0., 0., 0., 0., 0., 0., 0., 0., 0., 0., 0., 0., 0., 0., 0., 0., 0., 0.,
        0., 0., 0., 0., 0., 0., 0., 0., 0., 0., 0., 0., 0., 0., 0., 0., 0., 0.,
        0., 0., 0., 0., 0., 0., 0., 0., 0., 0., 0., 0., 0., 0., 0., 0., 0., 0.,
        0., 0., 0., 0., 0., 0., 0., 0., 0., 0., 0., 0., 0., 0., 0., 0., 0., 0.,
        0., 0., 0., 0., 0., 0., 0., 0., 0., 0., 0., 0., 0., 0., 0., 0., 0., 0.,
        0., 0., 0., 0., 0., 0., 0., 0., 0., 0., 0., 0., 0., 0., 0., 0., 0., 0.,
        0., 0., 0., 0., 0., 0., 0., 0., 0., 0., 0., 0., 0., 0., 0., 0., 0., 0.,
        0., 0., 0., 0., 0., 0., 0., 0., 0., 0., 0., 0., 0., 0., 0., 0., 0., 0.,
        0., 0., 0., 0., 0., 0., 0., 0., 0., 0., 0., 0., 0., 0., 0., 0., 0., 0.,
        0., 0., 0., 0., 0., 0., 0., 0., 0., 0., 0., 0., 0., 0., 0., 0., 0., 0.,
        0.], dtype=torch.float64)
```

[6]:
```
len(one_hot_encoding)
```

[6]:
```
343
```

[7]:
```
one_hot_encoding["CD3E"].argmax()
```

[7]: tensor(52)

[8]:
```
one_hot_encoding_int = dict()
for key in one_hot_encoding.keys():
    one_hot_encoding_int[key] = one_hot_encoding[key].argmax()
```

[9]:
```
one_hot_encoding_int["CD3E"]
```

[9]: tensor(52)

##### 1.1 Generate sample graph

Since the sample size is not too large, the entire sample area is processed without tiling, i.e., one tile per sample. The spatial RNA graph is constructed with radius 3.0. Setting the `process_mode` as "tile" will generate the tile graph data, and setting `process_mode` as "subgraph" will generate subgraph from the tile graph data.

[10]:
```
SpatialRNA(
    root= "../data/"+f'{sample_name}'+"/",
    sample_name=f'{sample_name}',
    one_hot_encoding=one_hot_encoding_int,
    num_tiles=1,
    dim_x = "x_location",
    dim_y = "y_location",
    tile_by_dim="y_location",
    process_mode="tile",
    load_type="blank",
    feature_col="feature_name",
    num_neighbours=[-1,-1],
    # num_seed_nodes=5000,
    # num_walks=5,
    radius_r=3.0

)
```

```
None  not exist, nothing loaded
```

[10]: SpatialRNA()

[11]:
```
%ls ../data/VUHD038/
```

```
processed/  raw/  subgraph/
```

[12]:
```
%ls ../data/VUHD038/processed
```

```
pre_filter.pt  pre_transform.pt  VUHD038_data_tile0.pt
```

##### 1.2 Generate a subgraph in a node-based manner with random walks

The subgraph construction starts with seed nodes sampling (5,000 seed nodes), and random walks are performed starting from the seed nodes with 5 walks per node. The nodes reaches by these random walks are included in the subgraph including their 2-hop neighbours.

[13]:
```
sub_g = SpatialRNA(
    root= "../data/"+f'{sample_name}'+"/",
    sample_name=f'{sample_name}',
    one_hot_encoding=one_hot_encoding_int,
    num_tiles=1,
    radius_r=3.0,
    dim_x = "x_location",
    dim_y = "y_location",
    tile_by_dim="y_location",
    load_type="subgraph",
    feature_col="feature_name",
    process_mode="subgraph",
    subgraph_mode="node_based",
    num_seed_nodes=5000,
    # batch_size is inferred
    num_walks=5,
    num_neighbors=[-1,-1],
    force_resample=False,
    subgraph_type="bidirectional"
)
```

```
To generate subgraphs for
['../data/VUHD038/subgraph/VUHD038_subgraph_data_tile0.pt']
Subgraphs already exist, skipping subgraph generation
loading from file  ../data/VUHD038/subgraph/VUHD038_subgraph_data_tile0.pt
```

[14]:
```
## loaded data is a list of size 1
sub_g[0]
```

[14]:
```
Data(x=[477476], edge_index=[2, 20038542], edge_label_index=[2, 25000])
```

[18]:
```
%ls ../data/VUHD038/subgraph/
```

```
VUHD038_subgraph_data_tile0.pt
```

##### 1.3 Repeat for all 45 samples to create the subgraphs

Repeat the subgraph generation process for the 45 samples like above.

#### 2 Join subgraphs for model training

Once subgraphs from 45 samples are ready, we can join them and use the joined graph for GNN training.

[19]:
```
sample_names =
    [
        "THD0008","TILD080LA","TILD117MA1","TILD175MA","VUHD049",
        "VUHD113","VUILD102MA","VUILD105MA2","VUILD115MA","VUILD48LA2",
        "VUILD78MA","VUILD96MA","THD0011","TILD111LA","TILD117MA2",
        "TILD299MA","VUHD069","VUHD116A","VUILD104MA1","VUILD106MA",
        "VUILD141MA","VUILD49LA","VUILD91LA","TILD028LA","TILD113LA",
        "TILD130LA","TILD315MA","VUHD090","VUHD116B","VUILD104MA2",
        "VUILD107MA","VUILD142MA","VUILD58MA","VUILD91MA","TILD049MA",
        "TILD117LA","TILD167LA","VUHD038","VUHD095","VUILD102LA",
        "VUILD105MA1","VUILD110LA","VUILD48LA1","VUILD78LA","VUILD96LA"
    ]
```

[20]:
```
subgraph_list  = ["../data/"+f"{sample_name}/subgraph/
 ↪"+f"{sample_name}_subgraph_data_tile0.pt" for sample_name in sample_names]
subgraph_list
```

[20]:
```
['../data/THD0008/subgraph/THD0008_subgraph_data_tile0.pt',
 '../data/TILD080LA/subgraph/TILD080LA_subgraph_data_tile0.pt',
 '../data/TILD117MA1/subgraph/TILD117MA1_subgraph_data_tile0.pt',
 '../data/TILD175MA/subgraph/TILD175MA_subgraph_data_tile0.pt',
 '../data/VUHD049/subgraph/VUHD049_subgraph_data_tile0.pt',
 '../data/VUHD113/subgraph/VUHD113_subgraph_data_tile0.pt',
 '../data/VUILD102MA/subgraph/VUILD102MA_subgraph_data_tile0.pt',
 '../data/VUILD105MA2/subgraph/VUILD105MA2_subgraph_data_tile0.pt',
 '../data/VUILD115MA/subgraph/VUILD115MA_subgraph_data_tile0.pt',
 '../data/VUILD48LA2/subgraph/VUILD48LA2_subgraph_data_tile0.pt',
 '../data/VUILD78MA/subgraph/VUILD78MA_subgraph_data_tile0.pt',
 '../data/VUILD96MA/subgraph/VUILD96MA_subgraph_data_tile0.pt',
 '../data/THD0011/subgraph/THD0011_subgraph_data_tile0.pt',
 '../data/TILD111LA/subgraph/TILD111LA_subgraph_data_tile0.pt',
 '../data/TILD117MA2/subgraph/TILD117MA2_subgraph_data_tile0.pt',
 '../data/TILD299MA/subgraph/TILD299MA_subgraph_data_tile0.pt',
 '../data/VUHD069/subgraph/VUHD069_subgraph_data_tile0.pt',
 '../data/VUHD116A/subgraph/VUHD116A_subgraph_data_tile0.pt',
 '../data/VUILD104MA1/subgraph/VUILD104MA1_subgraph_data_tile0.pt',
 '../data/VUILD106MA/subgraph/VUILD106MA_subgraph_data_tile0.pt',
 '../data/VUILD141MA/subgraph/VUILD141MA_subgraph_data_tile0.pt',
 '../data/VUILD49LA/subgraph/VUILD49LA_subgraph_data_tile0.pt',
 '../data/VUILD91LA/subgraph/VUILD91LA_subgraph_data_tile0.pt',
 '../data/TILD028LA/subgraph/TILD028LA_subgraph_data_tile0.pt',
 '../data/TILD113LA/subgraph/TILD113LA_subgraph_data_tile0.pt',
 '../data/TILD130LA/subgraph/TILD130LA_subgraph_data_tile0.pt',
 '../data/TILD315MA/subgraph/TILD315MA_subgraph_data_tile0.pt',
 '../data/VUHD090/subgraph/VUHD090_subgraph_data_tile0.pt',
 '../data/VUHD116B/subgraph/VUHD116B_subgraph_data_tile0.pt',
 '../data/VUILD104MA2/subgraph/VUILD104MA2_subgraph_data_tile0.pt',
```

```
 '../data/VUILD107MA/subgraph/VUILD107MA_subgraph_data_tile0.pt',
 '../data/VUILD142MA/subgraph/VUILD142MA_subgraph_data_tile0.pt',
 '../data/VUILD58MA/subgraph/VUILD58MA_subgraph_data_tile0.pt',
 '../data/VUILD91MA/subgraph/VUILD91MA_subgraph_data_tile0.pt',
 '../data/TILD049MA/subgraph/TILD049MA_subgraph_data_tile0.pt',
 '../data/TILD117LA/subgraph/TILD117LA_subgraph_data_tile0.pt',
 '../data/TILD167LA/subgraph/TILD167LA_subgraph_data_tile0.pt',
 '../data/VUHD038/subgraph/VUHD038_subgraph_data_tile0.pt',
 '../data/VUHD095/subgraph/VUHD095_subgraph_data_tile0.pt',
 '../data/VUILD102LA/subgraph/VUILD102LA_subgraph_data_tile0.pt',
 '../data/VUILD105MA1/subgraph/VUILD105MA1_subgraph_data_tile0.pt',
 '../data/VUILD110LA/subgraph/VUILD110LA_subgraph_data_tile0.pt',
 '../data/VUILD48LA1/subgraph/VUILD48LA1_subgraph_data_tile0.pt',
 '../data/VUILD78LA/subgraph/VUILD78LA_subgraph_data_tile0.pt',
 '../data/VUILD96LA/subgraph/VUILD96LA_subgraph_data_tile0.pt']
```

[21]:
```
from torch_geometric.data import Batch,Data
```

[22]:
```
data_list = [Data(**(torch.load(data_path,weights_only=True)[0])) for data_path␣
 ↪in subgraph_list]
```

[23]:
```
len(data_list)
```

[23]:
```
45
```

[24]:
```
data_list[0]
```

[24]:
```
Data(x=[1264610], edge_index=[2, 70938630], edge_label_index=[2, 25000])
```

#### 2.1 Create BatchData from merging all subgraphs

[ ]:
```
d_batch = Batch.from_data_list(data_list)
d_batch.x.shape
```

[26]:
```
d_batch
```

[26]:
```
DataBatch(x=[42447051], edge_index=[2, 2015025088], edge_label_index=[2,
1125000], batch=[42447051], ptr=[46])
```

[27]:
```
d_batch.ptr
```

[27]:
```
tensor([       0,  1264610,  2490908,  3634647,  4879671,  5740954,  6597488,
         7672922,  8456355,  9318554, 10315050, 10928126, 11994914, 12793121,
        13727665, 14961873, 16363320, 17218194, 18197059, 19144352, 20140282,
        20767010, 21835574, 22852247, 23843680, 24702689, 25693713, 26925009,
        27613591, 28584626, 29495681, 30645948, 30890595, 31627143, 32687055,
        33530880, 34705755, 35297115, 35774591, 36491949, 37443865, 38376891,
        39642066, 40605009, 41355417, 42447051])
```

[28]:
```
import random as rn
rn.seed(1024)

device = torch.device('cuda' if torch.cuda.is_available() else 'cpu')

print(torch.cuda.is_available())
print(device)
```

```
True
cuda
```

[29]:
```
from torch_geometric.nn import GATConv

class GATWithL2Normalization(torch.nn.Module):
    def __init__(self, in_channels, hidden_channels, out_channels, num_heads=1,␣
 ↪dropout=0.0):
        super(GATWithL2Normalization, self).__init__()
        self.conv1 = GATConv(in_channels, hidden_channels, heads=num_heads,␣
 ↪dropout=dropout)
        self.conv2 = GATConv(hidden_channels * num_heads, out_channels, heads=1,␣
 ↪concat=False, dropout=dropout)

    def forward(self, x, edge_index):
        # First GAT layer
        x = self.conv1(x, edge_index)
        x = F.elu(x)
        x = F.normalize(x, p=2, dim=1)  # L2-normalize the embeddings

        # Second GAT layer
        x = self.conv2(x, edge_index)
        x = F.normalize(x, p=2, dim=1)  # L2-normalize again after the second␣
 ↪layer

        return x

model = GATWithL2Normalization(
   in_channels = 343,
   hidden_channels=50,
   out_channels = 50
).to(device)
```

[30]:
```
## Easiy swop the models, for example

# model = GAT(
#       343,
#       hidden_channels=50,
#       num_layers=2,
```

```
#       v2 = use_v2,
#   ).to(device)
```

[31]:
```
print(model)
print("d_batch.x.size() ", d_batch.x.size())
print("d_batch.edge_index.size() ", d_batch.edge_index.size())

print("d_batch.edge_label_index.size() ", d_batch.edge_label_index.size())
```

```
GATWithL2Normalization(
  (conv1): GATConv(343, 50, heads=1)
  (conv2): GATConv(50, 50, heads=1)
)
d_batch.x.size()  torch.Size([42447051])
d_batch.edge_index.size()  torch.Size([2, 2015025088])
d_batch.edge_label_index.size()  torch.Size([2, 1125000])
```

#### 2.2 Set up training data loader from the joined graph

[32]:
```
from torch_geometric.loader import LinkNeighborLoader
from torch_geometric.sampler import NegativeSampling

train_loader = LinkNeighborLoader(
    d_batch,
    batch_size=2048,
    shuffle=True,
    edge_label_index = d_batch.edge_label_index,
    neg_sampling=NegativeSampling(mode="triplet",amount=1),
    # weight breaks sampling due to large number of nodes.
    ␣
 ↪#neg_sampling=NegativeSampling(mode="triplet",amount=1,dst_weight=degree(d_batch.
 ↪edge_index[0])**0.75),
    num_neighbors=[20,10],
    disjoint = False,
    subgraph_type = "bidirectional" ,
)
```

[33]:
```
optimizer = torch.optim.Adam(model.parameters(), lr=0.001)
def train():
    model.train()

    total_loss = 0
    batch_accuracy = []
    for batch in tqdm(train_loader):
        batch = batch.to(device)
        optimizer.zero_grad()
```

```
        #h = model(F.one_hot(batch.x, num_classes=gene_list.shape[0]).type(torch.
↪float), batch.edge_index)
        b_edge_label_index = torch.concat([torch.stack([batch.src_index, batch.
↪dst_pos_index]),
                                           torch.stack([batch.src_index, batch.
↪dst_neg_index])],1)
        b_edge_label = torch.concat([torch.tensor([1]*batch.src_index.
↪shape[0]),torch.tensor([0]*batch.src_index.shape[0])])

        batch.x = F.one_hot(batch.x, num_classes=343).float().squeeze(1)
        h = model(batch.x, batch.edge_index)
        h_src = h[b_edge_label_index[0]]
        h_dst = h[b_edge_label_index[1]]
        pred = (h_src * h_dst).sum(dim=-1)
        loss = F.binary_cross_entropy_with_logits(pred.to(device), b_edge_label.
↪float().to(device))
        # Step 1: Apply Sigmoid to convert logits to probabilities
        probabilities = torch.sigmoid(pred)
        # Step 2: Threshold probabilities to get binary predictions
        predictions = (probabilities >= 0.5).float()

        # Step 3: Calculate accuracy
        correct_predictions = (predictions == b_edge_label.float().to(device)).
↪float()
        batch_accuracy = [correct_predictions.sum().cpu() / pred.size(0)] +␣
↪batch_accuracy
        loss.backward()

        optimizer.step()
        total_loss += float(loss) * pred.size(0)
    return total_loss / d_batch.num_nodes, np.mean(batch_accuracy)
```

#### 2.3 Perform training and save the trained model

[34]:
```
import time
from tqdm import tqdm

times = []
train_loss = []
train_acc_list = []
for epoch in range(1, 6):
    start = time.time()
    loss,acc = train()
    print(f'Epoch: {epoch:03d}, Loss: {loss:.4f}, Train_Acc: {acc:.2f}')
    times.append(time.time() - start)
    train_loss = train_loss + [loss]
```

```
    train_acc_list = train_acc_list + [acc]
print(f"Median time per epoch: {torch.tensor(times).median():.4f}s")
```

```
100%|██████████| 550/550 [02:58<00:00,  3.09it/s]

Epoch: 001, Loss: 0.0275, Train_Acc: 0.79

100%|██████████| 550/550 [02:55<00:00,  3.13it/s]

Epoch: 002, Loss: 0.0274, Train_Acc: 0.81

100%|██████████| 550/550 [02:48<00:00,  3.26it/s]

Epoch: 003, Loss: 0.0273, Train_Acc: 0.81

100%|██████████| 550/550 [02:49<00:00,  3.25it/s]

Epoch: 004, Loss: 0.0273, Train_Acc: 0.81

100%|██████████| 550/550 [02:50<00:00,  3.23it/s]

Epoch: 005, Loss: 0.0273, Train_Acc: 0.81
Median time per epoch: 170.1486s
```

[ ]:
```
model
```

[ ]:
```
out_model = "../output/trained_model/GATL2_trained.model_weights.pth"
torch.save(model,out_model)
```

### 3 Plot spatial tissue domains in lung samples

We run GMM clustering using all transcript embeddings using [../code/run_gmm.py], and we now aggregate the cluster outcomes in hex bins and plot them.

[5]:
```
import os
import pandas as pd
import numpy as np
import matplotlib.pyplot as plt
import seaborn as sns
from scipy.stats import mode
```

[45]:
```
# define a helper function for appending cluster result to sample meta
def join_meta(sample_name):
    sample_gmm_res = "../output/GATv2_trained/epoch5/gmm_clusters/gmm12_trained/
↪"+f"{sample_name}_noMeans.txt"
    sample_gmm_res = pd.read_csv(sample_gmm_res,names=["cluster_id"])
    sample_meta = pd.read_csv("../data/"+f"{sample_name}/raw/{sample_name}.
↪csv",usecols=["x_location","y_location"])
    sample_meta["cluster_id"] = sample_gmm_res.cluster_id
    return sample_meta
```

[21]:
```python
from matplotlib.colors import ListedColormap
```

[22]:
```python
ListedColormap(plt.cm.tab20.colors[:12])
```

[22]:

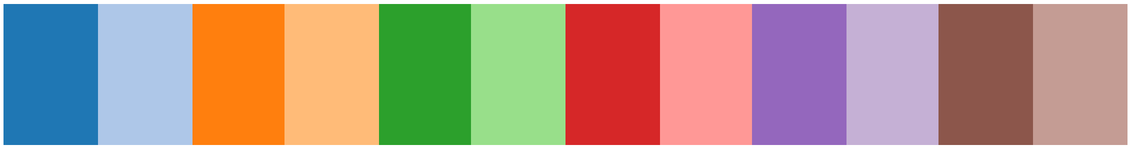

[27]:
```python
def get_major_cluster(values,bin_thresh=10):
    if len(values) < bin_thresh:
        return np.nan  # Return NaN for empty bins
    result = mode(values, keepdims=True)  # Ensure the result is always array-like
    return result.mode[0] if result.mode.size > 0 else np.nan

# Hex bin size
bin_width = 5
x_bins = int((sample_meta['x_location'].max() - sample_meta['x_location'].min()) / bin_width)
y_bins = int((sample_meta['y_location'].max() - sample_meta['y_location'].min()) / bin_width)
```

[58]:
```python
# Define the grid
fig, axes = plt.subplots(2, 2, figsize=(12, 10), constrained_layout=True)
axes = axes.flatten()

# Loop over the samples and axes
for ax, sample_name in zip(axes, ["VUHD116A","VUILD91LA","TILD080LA","VUHD049"]):
    sample_meta = join_meta(sample_name)  # Get the sample's metadata
    # Create the hexbin plot
    hb = ax.hexbin(
        sample_meta['x_location'],
        sample_meta['y_location'],
        C=sample_meta['cluster_id'],
        gridsize=(x_bins, y_bins),
        reduce_C_function=get_major_cluster,
        linewidths=0.05,
        cmap=ListedColormap(plt.cm.tab20.colors[:12])
    )
    # Customize the plot
    ax.invert_yaxis()
    ax.set_xlabel(sample_name, fontsize=12)
    ax.set_ylabel("Y Location", fontsize=12)
```

```
    ax.tick_params(axis='both', which='both', colors='black')
    ax.spines['top'].set_visible(False)
    ax.spines['right'].set_visible(False)
    ax.spines['left'].set_linewidth(1.5)
    ax.spines['bottom'].set_linewidth(1.5)
    ax.grid(False)

    # Add a colorbar for each subplot
    cb = fig.colorbar(hb, ax=ax, orientation='vertical')
    cb.set_label("Major Cluster", fontsize=10)

fig.suptitle("Hexbin Plots for Four Samples", fontsize=12, y=1.02)
fig.dpi = 300
plt.show()
```

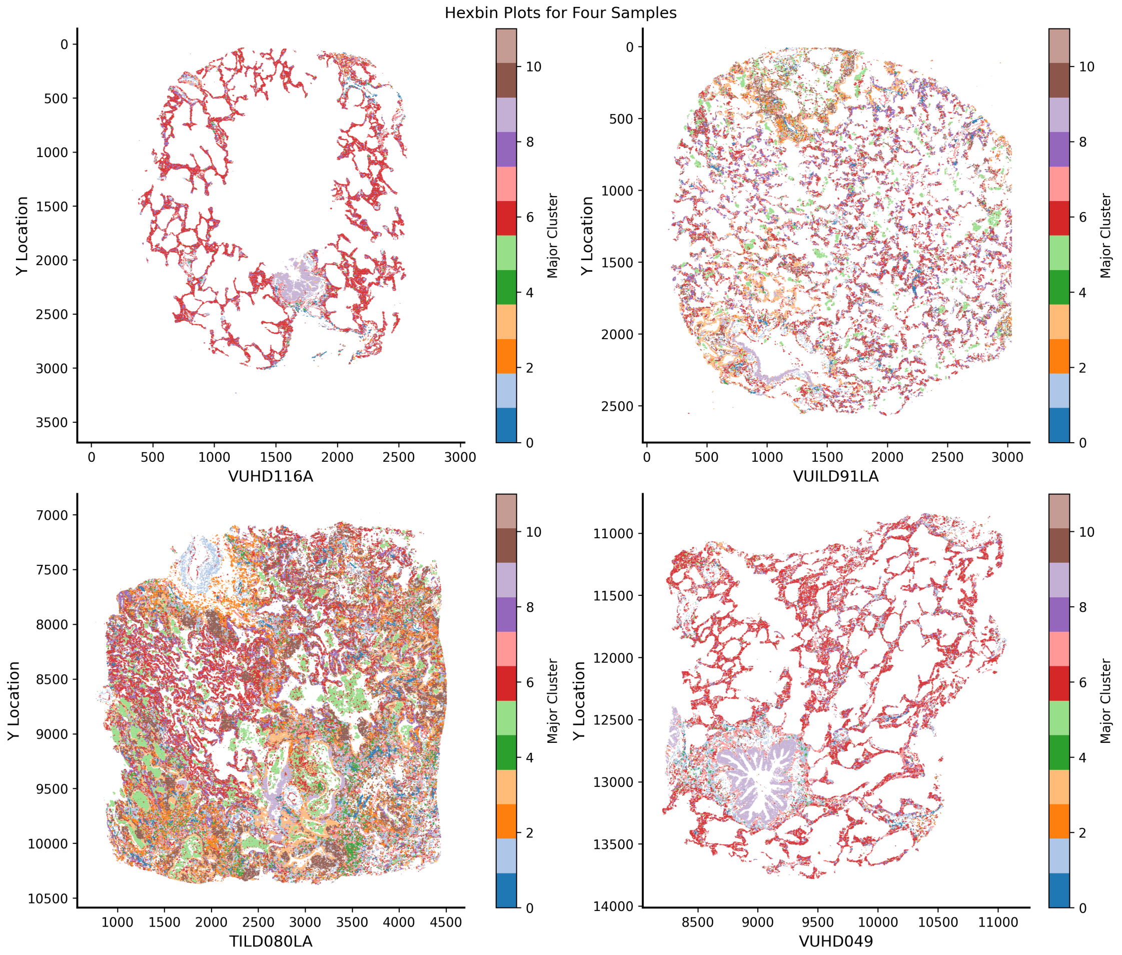
