## Supplementary_case_study2 for "SpatialRNA: a python package for easy application of Graph Neural Network models on single-molecule spatial transcriptomics dataset"

### case_study_xenium_5k_ovariancancer

March 19, 2025

**3 Join subgraphs for model training** **12**

**4 Inference of transcript embeddings with the trained model** **17**

**5 Generate hexbin plot visualisation** **19**

### 1 10x Xenium5k OvarianCancer

In this report, we will have an overview of the `Xenium5k_10x_OvarianCancer` dataset and prepare the input csv file for GNN anallysis.

The 10x_Xenium5k_OvarianCancer dataset is a high-resolution imaging-based spatial transcriptomics dataset generated using the 10x Genomics Xenium platform. It captures spatially resolved gene expression data from ovarian cancer tissue, offering insights into the cellular architecture and microenvironment of the tumor. This dataset includes approximately 5,000 unique transcripts per region, allowing detailed analysis of cell type distribution, gene expression patterns, and tumor-immune interactions within the ovarian cancer microenvironment.

We downloaded the provided data files from 10x Genomics website, and here we will focus on the transcript file which contains the spatially detected transcripts from 5,000 genes from the ovarian cancer tissue.

[1]:
```
import pandas as pd
import numpy as np
```

```
import sys
import torch
import torch.nn.functional as F
```

[ ]:
```
file_path = "/mnt/beegfs/mccarthy/backed_up/general/Datasets/
↪Xenium5k_10x_OvarianCancer/transcripts.parquet"
# Load the Parquet file into a pandas DataFrame
detected_tx = pd.read_parquet(file_path)
```

[6]:
```
detected_tx.shape
```

[6]:
```
(147706750, 13)
```

There are approximately 147.7 million transcripts were detected.

#### 1.1 Transcript filtering

We filter the list of transcripts by removing control probes as well as transcripts with `qv` $< 20$.

[7]:
```
detected_tx = detected_tx[detected_tx.qv >=20]
```

[8]:
```
detected_tx.shape
```

[8]:
```
(130166566, 13)
```

[9]:
```
detected_tx
```

[9]:
```
             transcript_id     cell_id  overlaps_nucleus  \
0          281681135150958  UNASSIGNED                 0
1          281681135148861  UNASSIGNED                 0
2          281681135142689  UNASSIGNED                 0
3          281681135151342  UNASSIGNED                 0
4          281681135145606  UNASSIGNED                 0
...                    ...         ...               ...
147706691  282213711085714  UNASSIGNED                 0
147706746  281569465992873  UNASSIGNED                 0
147706747  281569465992499  UNASSIGNED                 0
147706748  281569465993016  UNASSIGNED                 0
147706749  281569465992013  UNASSIGNED                 0

                     feature_name    x_location   y_location  z_location  \
0                            AAMP    158.312500  2545.921875   12.812500
1                             AXL    104.546875  2650.625000   13.078125
2                          CSNK2B    107.156250  2742.234375   13.234375
3                           GANAB    235.765625  2723.046875   13.218750
4                             IDE    155.906250  2544.421875   13.046875
...                           ...           ...          ...         ...
147706691  DeprecatedCodeword_17634  11391.640625  7003.203125   28.046875
```

```
147706746                        DDX43  11263.296875  802.640625  23.171875
147706747                        FOXO1  11252.703125  799.203125  23.015625
147706748                         OCRL  11301.562500  910.703125  23.140625
147706749                       ZNF683  11462.656250  856.390625  22.937500

              qv fov_name  nucleus_distance  codeword_index  \
0          40.00       D1        209.296875            2394
1          40.00       D1        224.453125           15901
2          32.25       D1        233.234375           13794
3          21.25       D1        107.578125           15280
4          26.25       D1        212.140625            1823
...          ...      ...               ...             ...
147706691  21.25      J19        454.750000           17634
147706746  37.50      B19        622.750000            3947
147706747  37.25      B19        619.312500           13588
147706748  40.00      B19        560.187500           11793
147706749  40.00      B19        682.218750           16219

             codeword_category  is_gene
0              predesigned_gene     True
1              predesigned_gene     True
2              predesigned_gene     True
3              predesigned_gene     True
4              predesigned_gene     True
...                         ...      ...
147706691  deprecated_codeword    False
147706746     predesigned_gene     True
147706747     predesigned_gene     True
147706748     predesigned_gene     True
147706749     predesigned_gene     True

[130166566 rows x 13 columns]
```

[10]:
```
np.unique(detected_tx.codeword_category)
```

[10]:
```
array(['custom_gene', 'deprecated_codeword', 'genomic_control_probe',
       'negative_control_codeword', 'negative_control_probe',
       'predesigned_gene', 'unassigned_codeword'], dtype=object)
```

[11]:
```
detected_tx = detected_tx[detected_tx.is_gene]
```

[12]:
```
detected_tx.shape
```

[12]:
```
(120455566, 13)
```

For this case study, we also remove transcripts that were not included in cell boundaries.

[13]:
```
detected_tx = detected_tx[detected_tx.cell_id !="UNASSIGNED"]
```

[14]:
```
detected_tx.shape
```

[14]:
```
(106483418, 13)
```

#### 1.2 Gene panel

[15]:
```
gene_list = np.unique(detected_tx.feature_name)
```

[16]:
```
gene_list.shape
```

[16]:
```
(5101,)
```

[19]:
```
gene_list
```

[19]:
```
array(['A2ML1', 'AAMP', 'AAR2', ..., 'ZUP1', 'ZYG11B', 'ZYX'],
      dtype=object)
```

[27]:
```
x = torch.tensor(np.arange(gene_list.shape[0]))

one_hot_encoding = dict(zip(gene_list, F.one_hot(x, num_classes=gene_list.
 ↪shape[0]).type(torch.float)))
gene_to_int = {key: torch.argmax(value).item() for key, value in␣
 ↪one_hot_encoding.items()}
#gene_to_int
```

[38]:
```
# The full list of genes:
# pd.DataFrame({"feature_name":gene_to_int.keys(),"Int_ID":gene_to_int.
 ↪values()}).to_csv("./gene_list.csv")
```

[41]:
```
gene_count = detected_tx.feature_name.value_counts()
```

[43]:
```
np.mean(gene_count)
```

[43]:
```
np.float64(20875.008429719663)
```

#### 1.3 Gene panel filtering

We next filter the gene panel and remove genes that have very low counts.

[51]:
```
gene_count[gene_count > 500]
```

[51]:
```
feature_name
EEF1G     3513254
H19       1792472
CD47       592996
H3F3B      514389
HNRNPD     453642
            ...
```

```
NKX2-3        505
NOS1          504
PCDHA2        504
GATA1         503
POU4F3        501
Name: count, Length: 4925, dtype: int64
```

[55]:
```
filtered_gene_list = gene_count[gene_count > 500].index
```

[58]:
```
# Save the filtered gene panel
# pd.DataFrame(filtered_gene_list).to_csv("filtered_gene_list_min500.csv")
```

[59]:
```
detected_tx = detected_tx[detected_tx.feature_name.isin(filtered_gene_list)]
```

We now only keep transcripts originating from the filtered gene panel.

[62]:
```
(detected_tx).shape
```

[62]:
```
(106423555, 13)
```

[63]:
```
detected_tx
```

[63]:
```
              transcript_id     cell_id  overlaps_nucleus feature_name  \
94          281844344737011  ahflbdfk-1                 1         AAMP
95          281844346167904  ahflbdfk-1                 1         AAMP
96          281844345420828  ahfnbpmi-1                 1         AAMP
97          281844346045776  ahflbdfk-1                 1        ABCC2
98          281844344758312  ahfnbpmi-1                 1         ABL2
...                     ...         ...               ...          ...
147705883   282132106710640  nnifhkho-1                 0       ZFAND5
147705890   282050502453723  nniganjg-1                 0       ZNF768
147705891   282050502419023  nnfbhnnf-1                 1         ZUP1
147705893   282050502466715  nniceino-1                 1          ZYX
147705894   282050502418751  nniceino-1                 1          ZYX

              x_location   y_location  z_location     qv fov_name  \
94            244.562500  3744.984375   16.078125  35.75       F1
95            247.546875  3747.718750   12.562500  40.00       F1
96            248.875000  3738.406250   14.062500  40.00       F1
97            243.812500  3749.140625   17.515625  37.00       F1
98            249.968750  3736.515625   13.890625  40.00       F1
...                  ...          ...         ...    ...      ...
147705883   11336.250000  5821.484375   33.515625  22.00      I19
147705890   11319.484375  5809.187500   25.687500  40.00      H19
147705891   11382.750000  5758.968750   22.937500  40.00      H19
147705893   11251.562500  5828.046875   28.156250  40.00      H19
147705894   11253.640625  5828.328125   25.859375  40.00      H19
```

```
           nucleus_distance  codeword_index codeword_category  is_gene
94                 0.000000            2394  predesigned_gene     True
95                 0.000000            2394  predesigned_gene     True
96                 0.000000            2394  predesigned_gene     True
97                 0.000000           10866  predesigned_gene     True
98                 0.000000            8644  predesigned_gene     True
...                     ...             ...               ...      ...
147705883          0.578125            6489  predesigned_gene     True
147705890          4.125000            2931  predesigned_gene     True
147705891          0.000000           10346  predesigned_gene     True
147705893          0.000000           13575  predesigned_gene     True
147705894          0.000000           18454  predesigned_gene     True

[106423555 rows x 13 columns]
```

[64]:
```
(detected_tx.overlaps_nucleus).sum()
```

[64]:
```
np.uint64(55753917)
```

##### 1.4 Save to output

Save the final list of transcripts for GNN analysis to a csv file for GNN analysis.

[65]:
```
detected_tx.to_csv("../data/OvarianCancer/raw/OvarianCancer.csv")
```

#### 2 Tissue tiling and graph, subgraph generation

We use `SpatialRNA` to construct the data inputs for GNN training. We first tile the tissue along Y axis into 30 strips, and we build RNA spatial graphs for all tiles. Finally for each tile, we sample a subgraph, and subgraphs will be joined for training GNN models later.

[1]:
```
import pandas as pd
import numpy as np
from torch_geometric.nn import radius_graph
from torch_geometric import seed_everything
import torch
import os.path as osp
import time

import torch_geometric.transforms as T
from torch_geometric.loader import LinkNeighborLoader,NeighborLoader
from torch_geometric.nn import GraphSAGE,GAT
import torch.nn.functional as F
from torch_geometric.data import Data
from torch_geometric.transforms import RandomNodeSplit
import random
import sys
```

```
import matplotlib.pyplot as plt
from spatialrna import SpatialRNA
```

[2]:
```
gene_panel = pd.read_csv(
    "./filtered_gene_list_min500.csv")
gene_panel

gene_list = gene_panel.feature_name
x = torch.tensor(np.arange(gene_list.shape[0]))

one_hot_encoding = dict(zip(gene_list, F.one_hot(x, num_classes=gene_list.
 ↪shape[0]).type(torch.float)))
gene_to_int = {key: torch.argmax(value).item() for key, value in␣
 ↪one_hot_encoding.items()}
```

We convert the gene labels of transcripts into integer labels.

[3]:
```
gene_to_int["EEF1G"],one_hot_encoding["EEF1G"]
```

[3]:
```
(0, tensor([1., 0., 0.,  ..., 0., 0., 0.]))
```

#### 2.1 Create tissue tiles and tile graphs

`SpatialRNA` tiles the tissues according to the `num_tiles` supplied. The `process_tile_ids` controls which tile(s) to process for in one function call, and one tile at a time is recommended for large tissues. Processing tiles can be paralellised. Once all the tiles are processed, the spatil RNA graphs created from tiles are saved to the "processed" directory.

We can process all tiles in paralle as shown in the workflow ../workflows/run_generate_subg.smk

[4]:
```
%ls ../data/OvarianCancer
```

```
processed/  raw/  subgraph/
```

[5]:
```
?SpatialRNA
```

```
Init signature:
SpatialRNA(
    root: str,
    sample_name: Optional[str] = None,
    radius_r: float = 3.0,
    dim_x: str = 'X',
    dim_y: str = 'Y',
    tile_by_dim: str = 'Y',
    num_tiles: int = 1,
    process_mode: str = 'tile',
    load_type: str = 'tile',
    load_tile_id: int = 0,
    pad_hops: float = 2,
    feature_col='gene',
```

```
    max_num_neighbors: int = 500,
    process_tile_ids: List[int] = None,
    one_hot_encoding: dict = None,
    subgraph_mode: str = 'link_based',
    num_sampled_edges: int = 5000,
    num_seed_nodes: int = 5000,
    num_walks: int = 5,
    transform=None,
    pre_transform=None,
    force_reload: bool = False,
    force_resample: bool = False,
    log: bool = True,
    seed: int = 100,
    **kwargs,
) -> None
Docstring:
A spatial RNA graph dataset. One dataset per tissue sample, and optionally creates
tiles from a tissue area. For each generated data.pt, it contains the nodes in␣
 ↪core areas for
making inference for transcripts with complete neighbourhoods.

Args:
    root (str): Root directory where the dataset should be saved.
    sample_name (str): The name of this sample. Appended to the processed␣
 ↪[sample_name]_data.pt.
                        Using basename of root dir if not supplied.
    radius_r (float): The radius for building RNA graphs.
    tile_by_dim (str): Along which dimension to make tiles. It should be a column␣
 ↪in the transcript data.frame.
    num_tiles (int): The number of tiles to create from this tissue sample.
    process_mode (str): The mode of process can be either "tile" or "subgraph".
    load_type (str): If :obj:`"tile"`, load the tile graph data.
        If :obj:`"subgraph"`, load the tile subgraph data.
    load_tile_id (int): Select one tile to load.
    process_tile_ids (List[int]): Select tiles to process including generating␣
 ↪tile graph and subgraph.
    one_hot_encoding (dict): Dictionary that mapps gene names to integer (or␣
 ↪one-hot-encoding tensors).
        It is essential to apply the same dictionary for all samples that are␣
 ↪analyzed together.
    transform (callable, optional): A function/transform that takes in an
        :obj:`torch_geometric.data.Data` object and returns a transformed
        version. The data object will be transformed before every access.
        (default: :obj:`None`)
    pre_transform (callable, optional): A function/transform that takes in
        an :obj:`torch_geometric.data.Data` object and returns a
```

```
        transformed version. The data object will be transformed before
        being saved to disk. (default: :obj:`None`)
    force_reload (bool, optional): Whether to re-process the dataset.
        (default: :obj:`False`)
    force_resample (bool, optional): Whether to re-process the tile dataset to␣
↪generate subgraphs.
        (default: :obj:`False`)
    subgraph_mode (str): The mode of subgraph generation can be either␣
↪"link_based" or "node_based".
        (default: :obj:`link_based`)
    num_sampled_edges (int): sample this many edge_index as positive edges when␣
↪generating subgraph from a tile (used in LinkNeighborLoader).
    num_seed_nodes:(int, optional): Number of seed nodes for generating subgraphs.
        (default: :obj:`5000`)
    num_walks:(int, optional): Number of walks for generating subgraphs.
        (default: :obj:`5`)

    log (bool, optional): Whether to print any console output while
        downloading and processing the dataset. (default: :obj:`True`)
    **kwargs (optional): Additional arguments of
        :class:`torch.utils.data.LinkNeighborLoader`, such as :obj:`num_neighbors`
File:           /mnt/beegfs/mccarthy/general/backed_up/rlyu/Projects/spatialrna/
↪spatialrna/spatialrna.py
Type:           type
Subclasses:
```

We now load one processed tile data using the function below, and check out the loaded data object.

[6]:
```
tile28_graph = SpatialRNA(
    root ="../data/OvarianCancer/",
    one_hot_encoding = gene_to_int,
    sample_name = "OvarianCancer",
    force_reload=False, # whether to force reprocessing of the raw csv file to␣
↪generate the tile graph data.
    radius_r=3.0,
    num_tiles= 30,
    feature_col = "feature_name",
    dim_x="x_location",
    dim_y="y_location",
    tile_by_dim="y_location",
    load_tile_id=28,
    log=True,
    load_type="tile")
```

```
loading from file  ../data/OvarianCancer/processed/OvarianCancer_data_tile28.pt
```

[7]:
```
tile28_graph[0]
```

```
[7]: Data(x=[336775], edge_index=[2, 17187528], trans_id=[336775],
core_mask=[336775])
```

`trans_id` corresponds to the index ID of transcripts in the raw/OvarianCancer.csv file. `core_mask` indicates whether a transcript is in the core area or the padding area of the tile. We have a total of 30 tiles in the processed folder.

```
[8]: %ls ../data/OvarianCancer/processed/
```

```
OvarianCancer_data_tile0.pt   OvarianCancer_data_tile24.pt
OvarianCancer_data_tile10.pt  OvarianCancer_data_tile25.pt
OvarianCancer_data_tile11.pt  OvarianCancer_data_tile26.pt
OvarianCancer_data_tile12.pt  OvarianCancer_data_tile27.pt
OvarianCancer_data_tile13.pt  OvarianCancer_data_tile28.pt
OvarianCancer_data_tile14.pt  OvarianCancer_data_tile29.pt
OvarianCancer_data_tile15.pt  OvarianCancer_data_tile2.pt
OvarianCancer_data_tile16.pt  OvarianCancer_data_tile3.pt
OvarianCancer_data_tile17.pt  OvarianCancer_data_tile4.pt
OvarianCancer_data_tile18.pt  OvarianCancer_data_tile5.pt
OvarianCancer_data_tile19.pt  OvarianCancer_data_tile6.pt
OvarianCancer_data_tile1.pt   OvarianCancer_data_tile7.pt
OvarianCancer_data_tile20.pt  OvarianCancer_data_tile8.pt
OvarianCancer_data_tile21.pt  OvarianCancer_data_tile9.pt
OvarianCancer_data_tile22.pt  pre_filter.pt
OvarianCancer_data_tile23.pt  pre_transform.pt
```

#### 2.2 Create tile subgraphs

Each subgraph was generated by sampling 5,000 edges and the two-hop neighbours of the sampled nodes. For example, for tile 28, the subgraph is generated by:

```
[ ]: SpatialRNA(
    root= "../data/OvarianCancer",
    sample_name='OvarianCancer',
    num_tiles=30,
    radius_r=3.0,
    dim_x = "x_location",
    dim_y = "y_location",
    tile_by_dim="y_location",
    load_type="subgraph",
    feature_col="feature_name",
    process_mode="subgraph",
    subgraph_mode="link_based",
    process_tile_ids=[28],
    num_sampled_edges = 5000,
    num_neighbors = [-1,-1], # all neighbours at first hop and the second hop
    force_resample=True,
    batch_size = 5000,
    log=True,
```

```
    subgraph_type="induced",
    shuffle=False
)
```

We have 30 subgraph data files in the subgraph folder.

[9]:
```
%ls ../data/OvarianCancer/subgraph/
```

```
OvarianCancer_subgraph_data_tile0.pt   OvarianCancer_subgraph_data_tile23.pt
OvarianCancer_subgraph_data_tile10.pt  OvarianCancer_subgraph_data_tile24.pt
OvarianCancer_subgraph_data_tile11.pt  OvarianCancer_subgraph_data_tile25.pt
OvarianCancer_subgraph_data_tile12.pt  OvarianCancer_subgraph_data_tile26.pt
OvarianCancer_subgraph_data_tile13.pt  OvarianCancer_subgraph_data_tile27.pt
OvarianCancer_subgraph_data_tile14.pt  OvarianCancer_subgraph_data_tile28.pt
OvarianCancer_subgraph_data_tile15.pt  OvarianCancer_subgraph_data_tile29.pt
OvarianCancer_subgraph_data_tile16.pt  OvarianCancer_subgraph_data_tile2.pt
OvarianCancer_subgraph_data_tile17.pt  OvarianCancer_subgraph_data_tile3.pt
OvarianCancer_subgraph_data_tile18.pt  OvarianCancer_subgraph_data_tile4.pt
OvarianCancer_subgraph_data_tile19.pt  OvarianCancer_subgraph_data_tile5.pt
OvarianCancer_subgraph_data_tile1.pt   OvarianCancer_subgraph_data_tile6.pt
OvarianCancer_subgraph_data_tile20.pt  OvarianCancer_subgraph_data_tile7.pt
OvarianCancer_subgraph_data_tile21.pt  OvarianCancer_subgraph_data_tile8.pt
OvarianCancer_subgraph_data_tile22.pt  OvarianCancer_subgraph_data_tile9.pt
```

[10]:
```
# check subgraph of tile 28
tile28_subgraph = SpatialRNA(
    root ="../data/OvarianCancer/",
    one_hot_encoding = gene_to_int,
    sample_name = "OvarianCancer",
    force_reload=False, # do not reprocess the raw file to generate the tile␣
↪graph data.
    radius_r=3.0,
    num_tiles= 30,
    feature_col = "feature_name",
    dim_x="x_location",
    dim_y="y_location",
    tile_by_dim="y_location",
    load_tile_id=28,
    log=True,
    load_type="subgraph")
```

```
loading from file
../data/OvarianCancer/subgraph/OvarianCancer_subgraph_data_tile28.pt
```

[11]:
```
tile28_subgraph[0]
```

[11]:
```
Data(x=[238295], edge_index=[2, 14348386], edge_label_index=[2, 5000])
```

In this data file, the x slot holds the input feature of each node and currently they are integer IDs representing gene labels. `edge_index` contains edges in the subgraph. `edge_label_index` contains the sampled 5,000 edges for subgraph construction, and we use these edges as the positive pairs in the model training step.

#### 3 Join subgraphs for model training

Join subgraphs from 15 tiles for model training.

[12]:
```
subgraph_list  = ["../data/OvarianCancer/subgraph/
↪OvarianCancer_subgraph_data_tile"+f'{e}.pt' for e in range(0,30,2)]
subgraph_list
```

[12]:
```
['../data/OvarianCancer/subgraph/OvarianCancer_subgraph_data_tile0.pt',
 '../data/OvarianCancer/subgraph/OvarianCancer_subgraph_data_tile2.pt',
 '../data/OvarianCancer/subgraph/OvarianCancer_subgraph_data_tile4.pt',
 '../data/OvarianCancer/subgraph/OvarianCancer_subgraph_data_tile6.pt',
 '../data/OvarianCancer/subgraph/OvarianCancer_subgraph_data_tile8.pt',
 '../data/OvarianCancer/subgraph/OvarianCancer_subgraph_data_tile10.pt',
 '../data/OvarianCancer/subgraph/OvarianCancer_subgraph_data_tile12.pt',
 '../data/OvarianCancer/subgraph/OvarianCancer_subgraph_data_tile14.pt',
 '../data/OvarianCancer/subgraph/OvarianCancer_subgraph_data_tile16.pt',
 '../data/OvarianCancer/subgraph/OvarianCancer_subgraph_data_tile18.pt',
 '../data/OvarianCancer/subgraph/OvarianCancer_subgraph_data_tile20.pt',
 '../data/OvarianCancer/subgraph/OvarianCancer_subgraph_data_tile22.pt',
 '../data/OvarianCancer/subgraph/OvarianCancer_subgraph_data_tile24.pt',
 '../data/OvarianCancer/subgraph/OvarianCancer_subgraph_data_tile26.pt',
 '../data/OvarianCancer/subgraph/OvarianCancer_subgraph_data_tile28.pt']
```

[ ]:
```
from torch_geometric.data import Batch,Data
data_list = [Data(**(torch.load(data_path,weights_only=True)[0])) for data_path␣
↪in subgraph_list]

len(data_list)
```

Create Batch data from the list of subgraphs

[ ]:
```
d_batch = Batch.from_data_list(data_list)

d_batch.x.shape
d_batch
```

[ ]:
```
device = torch.device('cuda' if torch.cuda.is_available() else 'cpu')

print(torch.cuda.is_available())
print(device)
```

[ ]:
```
from torch_geometric.loader import LinkNeighborLoader, NeighborLoader
from torch_geometric.nn import GraphSAGE, GAT, GATConv, AttentionalAggregation
```

```python
from torch_geometric.nn import GATConv

model_select = "GATL2"
```

[ ]:
```python
len(gene_to_int)
```

[ ]:
```python
input_feature_size = len(gene_to_int)
```

[ ]:
```python

```

[ ]:
```python
# construct the GNN model
class GATWithL2Normalization(torch.nn.Module):
    def __init__(self, in_channels, hidden_channels, out_channels, num_heads=1,␣
 ↪dropout=0.0):
        super(GATWithL2Normalization, self).__init__()
        self.conv1 = GATConv(in_channels, hidden_channels, heads=num_heads,␣
 ↪dropout=dropout)
        self.conv2 = GATConv(hidden_channels * num_heads, out_channels, heads=1,␣
 ↪concat=False, dropout=dropout)

    def forward(self, x, edge_index):
        # First GAT layer
        x = self.conv1(x, edge_index)
        x = F.elu(x)
        x = F.normalize(x, p=2, dim=1)  # L2-normalize the embeddings

        # Second GAT layer
        x = self.conv2(x, edge_index)
        x = F.normalize(x, p=2, dim=1)  # L2-normalize again after the second␣
 ↪layer

        return x
```

[ ]:
```python
if model_select == "GAT":
    model = GAT(
         input_feature_size,
         hidden_channels=50,
         num_layers=2,
         v2 = False,
     ).to(device)
elif model_select == "GATv2":
    model = GAT(
         input_feature_size,
         hidden_channels=100,
         num_layers=2,
         v2 = True,
     ).to(device)
```

```python
elif model_select == "GATL2":
    model = GATWithL2Normalization(
        in_channels = input_feature_size,
        hidden_channels=100,
        out_channels = 100
    ).to(device)
else:
    model = GraphSAGE(
         input_feature_size,
         hidden_channels=100,
         num_layers=2,
     ).to(device)

print(model)
print("d_batch.x.size() ", d_batch.x.size())
print("d_batch.edge_index.size() ", d_batch.edge_index.size())

print("d_batch.edge_label_index.size() ", d_batch.edge_label_index.size())
```

[ ]:
```python
from torch_geometric.loader import LinkNeighborLoader
from torch_geometric.sampler import NegativeSampling
from tqdm import tqdm
```

[ ]:
```python
train_loader = LinkNeighborLoader(
    d_batch,
    batch_size=200,
    shuffle=True,
    edge_label_index = d_batch.edge_label_index,
    neg_sampling=NegativeSampling(mode="triplet",amount=1),
    num_neighbors=[50,20],
    disjoint = False,
    subgraph_type = "bidirectional"

)
```

[ ]:
```python
optimizer = torch.optim.Adam(model.parameters(), lr=0.001)
def train():
    model.train()

    total_loss = 0
    batch_accuracy = []
    for batch in tqdm(train_loader):
        batch = batch.to(device)
        optimizer.zero_grad()
        b_edge_label_index = torch.concat([torch.stack([batch.src_index, batch.
 ↪dst_pos_index]),
```

```
                                            torch.stack([batch.src_index, batch.
↪dst_neg_index])],1)
        b_edge_label = torch.concat([torch.tensor([1]*batch.src_index.
↪shape[0]),torch.tensor([0]*batch.src_index.shape[0])])

        batch.x = F.one_hot(batch.x, num_classes=input_feature_size).float().
↪squeeze(1)
        h = model(batch.x, batch.edge_index)
        h_src = h[b_edge_label_index[0]]
        h_dst = h[b_edge_label_index[1]]
        pred = (h_src * h_dst).sum(dim=-1)
        loss = F.binary_cross_entropy_with_logits(pred.to(device), b_edge_label.
↪float().to(device))
        # Step 1: Apply Sigmoid to convert logits to probabilities
        probabilities = torch.sigmoid(pred)
        # Step 2: Threshold probabilities to get binary predictions
        predictions = (probabilities >= 0.5).float()

        # Step 3: Calculate accuracy
        correct_predictions = (predictions == b_edge_label.float().to(device)).
↪float()
        batch_accuracy = [correct_predictions.sum().cpu() / pred.size(0)] +␣
↪batch_accuracy
        loss.backward()

        optimizer.step()
        total_loss += float(loss) * pred.size(0)
    return total_loss / d_batch.num_nodes, np.mean(batch_accuracy)
```

[ ]:
```
epoch = 15
```

[ ]:
```
out_model = "../output/trained_model/"+model_select+"_trained/epoch"+f'{epoch}.
↪model_weights.pth'
```

[27]:
```
times = []
train_loss = []
train_acc_list = []
for epoch in range(1, epoch+1):
    start = time.time()
    loss,acc = train()
    #val_acc = val()
    #print(f'Epoch: {epoch:03d}, Loss: {loss:.4f}, Train_Acc: {acc:.2f}, Val_Acc:
↪ {val_acc:.2f}')
    print(f'Epoch: {epoch:03d}, Loss: {loss:.4f}, Train_Acc: {acc:.2f}')
    times.append(time.time() - start)
    train_loss = train_loss + [loss]
    train_acc_list = train_acc_list + [acc]
```

```python
print(f"Median time per epoch: {torch.tensor(times).median():.4f}s")
```

```
100%|██████████| 375/375 [01:35<00:00,  3.91it/s]

Epoch: 001, Loss: 0.0033, Train_Acc: 0.76

100%|██████████| 375/375 [01:29<00:00,  4.19it/s]

Epoch: 002, Loss: 0.0033, Train_Acc: 0.78

100%|██████████| 375/375 [01:28<00:00,  4.22it/s]

Epoch: 003, Loss: 0.0033, Train_Acc: 0.78

100%|██████████| 375/375 [01:24<00:00,  4.42it/s]

Epoch: 004, Loss: 0.0033, Train_Acc: 0.78

100%|██████████| 375/375 [01:24<00:00,  4.42it/s]

Epoch: 005, Loss: 0.0033, Train_Acc: 0.78

100%|██████████| 375/375 [01:24<00:00,  4.42it/s]

Epoch: 006, Loss: 0.0033, Train_Acc: 0.78

100%|██████████| 375/375 [01:27<00:00,  4.30it/s]

Epoch: 007, Loss: 0.0033, Train_Acc: 0.78

100%|██████████| 375/375 [01:59<00:00,  3.13it/s]

Epoch: 008, Loss: 0.0033, Train_Acc: 0.78

100%|██████████| 375/375 [02:12<00:00,  2.84it/s]

Epoch: 009, Loss: 0.0033, Train_Acc: 0.79

100%|██████████| 375/375 [01:24<00:00,  4.41it/s]

Epoch: 010, Loss: 0.0032, Train_Acc: 0.79

100%|██████████| 375/375 [01:24<00:00,  4.42it/s]

Epoch: 011, Loss: 0.0032, Train_Acc: 0.79

100%|██████████| 375/375 [01:24<00:00,  4.41it/s]

Epoch: 012, Loss: 0.0033, Train_Acc: 0.79

100%|██████████| 375/375 [01:24<00:00,  4.41it/s]

Epoch: 013, Loss: 0.0033, Train_Acc: 0.79

100%|██████████| 375/375 [01:24<00:00,  4.42it/s]

Epoch: 014, Loss: 0.0033, Train_Acc: 0.79

100%|██████████| 375/375 [01:25<00:00,  4.41it/s]

Epoch: 015, Loss: 0.0033, Train_Acc: 0.79
Median time per epoch: 84.9942s
```

[32]:
```
torch.save(model,out_model)
```

#### 4 Inference of transcript embeddings with the trained model

Inference of transcript embedding is performed per tile. Transcripts in the tissue tile have been labelled with core_mask, which is a binary marker that specify whether the transcripts are in the core area of the tile or come from the padding area. Only transcripts that are included in the core area are kept in the final output.

The following code chunks demonstrate the process of obtaining embeddings for transcripts in one tile.

[ ]:
```
import pandas as pd
import os.path as osp
import time
import sys

import torch
import torch.nn.functional as F
from torch_geometric.data import Data
from sklearn.linear_model import LogisticRegression

import torch_geometric.transforms as T
from torch_geometric.datasets import Planetoid
from torch_geometric.loader import LinkNeighborLoader, NeighborLoader
from torch_geometric.nn import GraphSAGE, GAT, GATConv, AttentionalAggregation
import numpy as np
import matplotlib.pyplot as plt
from torch_geometric.nn import radius_graph

import torch.nn.functional as F
from torch_geometric.nn import GATConv
from tqdm import tqdm
```

[2]:
```
class GATWithL2Normalization(torch.nn.Module):
    def __init__(self, in_channels, hidden_channels, out_channels, num_heads=1,␣
↪dropout=0.0):
        super(GATWithL2Normalization, self).__init__()
        self.conv1 = GATConv(in_channels, hidden_channels, heads=num_heads,␣
↪dropout=dropout)
        self.conv2 = GATConv(hidden_channels * num_heads, out_channels, heads=1,␣
↪concat=False, dropout=dropout)

    def forward(self, x, edge_index):
        # First GAT layer
```

```
        x = self.conv1(x, edge_index)
        x = F.elu(x)
        x = F.normalize(x, p=2, dim=1)  # L2-normalize the embeddings

        # Second GAT layer
        x = self.conv2(x, edge_index)
        x = F.normalize(x, p=2, dim=1)  # L2-normalize again after the second␣
↪layer

        return x
```

[7]:
```
model_select = "GATL2"
epoch = 15
out_model = "../output/trained_model/"+model_select+"_trained/epoch"+f'{epoch}.
↪model_weights.pth'
```

[8]:
```
## load the trained model
model = torch.load(out_model,weights_only=False)
model
```

[8]:
```
GATWithL2Normalization(
  (conv1): GATConv(4925, 100, heads=1)
  (conv2): GATConv(100, 100, heads=1)
)
```

[9]:
```
device = torch.device('cuda' if torch.cuda.is_available() else 'cpu')
```

[10]:
```
@torch.no_grad
def inference(subgraph_loader):
    xs = []
    for batch in tqdm(subgraph_loader):
        batch = batch.to(device)
        batch.x = torch.nn.functional.one_hot(batch.x, num_classes=4925).float().
↪squeeze(1)

        out = model(batch.x, batch.edge_index)
        # NOTE Only consider predictions and labels of seed nodes:
        # y = batch.y[:batch.batch_size]
        out = out[:batch.batch_size]
        xs.append(out.cpu())
    x_all = torch.cat(xs, dim=0)
    return x_all
```

[11]:
```
# Example of embedding prediction for molecules in one tile
```

[12]:
```
tile_embs = []
trans_ids = []
```

```
for tile_id in [0]:
    data_path = "../data/OvarianCancer/processed/
↪OvarianCancer_data_tile"+f'{tile_id}.pt'
    data = torch.load(data_path)
    data = Data(**data[0])
    subgraph_loader = NeighborLoader(
                            data,
                            input_nodes=data.core_mask,
                            #num_neighbors=[-1],
                            num_neighbors=[50,20],
                            batch_size=1024,
                            replace=False,
                            shuffle=False,
                            subgraph_type = "bidirectional"
                    )
    node_embs= inference(subgraph_loader).cpu()
    print(data.x.size())
    node_embs = node_embs.detach().numpy()
    # np.save("../output/"+f'{tile_id}.npy',node_embs)
    #tile_embs = tile_embs + [tile_emb]
    trans_ids = trans_ids + [data.trans_id[data.core_mask.cpu()]]
```

```
/tmp/ipykernel_536179/3423153253.py:5: FutureWarning: You are using `torch.load`
with `weights_only=False` (the current default value), which uses the default
pickle module implicitly. It is possible to construct malicious pickle data
which will execute arbitrary code during unpickling (See
https://github.com/pytorch/pytorch/blob/main/SECURITY.md#untrusted-models for
more details). In a future release, the default value for `weights_only` will be
flipped to `True`. This limits the functions that could be executed during
unpickling. Arbitrary objects will no longer be allowed to be loaded via this
mode unless they are explicitly allowlisted by the user via
`torch.serialization.add_safe_globals`. We recommend you start setting
`weights_only=True` for any use case where you don't have full control of the
loaded file. Please open an issue on GitHub for any issues related to this
experimental feature.
  data = torch.load(data_path)
100%|██████████| 1260/1260 [04:45<00:00,  4.42it/s]
```

```
torch.Size([1327724])
```

We use the `NeighborLoader` function to perform inference for batched input nodes, and we supplied the `input_nodes` with the `core_mask` of the data tile, therefore only nodes in the core area are inferred.

### 5 Generate hexbin plot visualisation

We now have obtained and clustered the transcripts embeddings for all transcripts detected in the input csv file. We now demonstrate a visualisation for examining the molecule-based spatial domain results.

[1]:
```python
import os
import pandas as pd
import numpy as np
import matplotlib.pyplot as plt
import seaborn as sns
from scipy.stats import mode
```

[ ]:
```python
## List all tiles
all_tiles = [f for f in os.listdir("../output/GATL2_trained/epoch10/gmm_clusters/
↪gmm12_trained/") if "tile" in f and f.endswith(".txt")]
all_tiles
```

We first obtain the transcript index ID of nodes in the data tiles so that we can map them to the transcripts with meta data from the input csv file.

[3]:
```python
# Aggregate transcript id files
all_tiles_tx_id = []
for tile_gmm in range(30):
    x = pd.read_csv(f"../output/GATL2_trained/epoch10/embeddings/
↪OvarianCancer_tile{tile_gmm}.csv")
    all_tiles_tx_id.append(x)
all_tiles_tx_id = pd.concat(all_tiles_tx_id, ignore_index=True)
```

[4]:
```python
all_tiles_tx_id
```

[4]:
```
               tx_id
0            8220009
1            8220010
2            8220011
3            8220012
4            8220013
...              ...
106423550  102176191
106423551  102176192
106423552  102176193
106423553  102176194
106423554  102176195

[106423555 rows x 1 columns]
```

[5]:
```python
print(all_tiles_tx_id.shape)
print(all_tiles_tx_id.head())
```

```
(106423555, 1)
     tx_id
0  8220009
1  8220010
```

```
2   8220011
3   8220012
4   8220013
```

[6]:
```
# Load all transcript meta data
all_tx = pd.read_csv("../data/OvarianCancer/raw/OvarianCancer.csv")
print(all_tx.shape)
```

```
(106423555, 14)
```

[7]:
```
# reorder based on `tx_id`
all_tx = all_tx.loc[all_tiles_tx_id['tx_id']].reset_index(drop=True)
print(all_tx.shape)
```

```
(106423555, 14)
```

##### 5.1 Load cluster IDs

We have clustered these transcripts based their embeddings using a python script, [../code/run_gmm.py]. We load the cluster IDs for all transcripts here.

[9]:
```
# Load cluster id
all_tiles_gmm12 = []
for tile_gmm in range(30):
    x = pd.read_csv(f"../output/GATL2_trained/epoch10/gmm_clusters/gmm12_trained/
↪OvarianCancer_tile{tile_gmm}.txt",
                    header=None, names=['gmm12'])
    all_tiles_gmm12.append(x)
```

[10]:
```
all_tiles_gmm12 = pd.concat(all_tiles_gmm12, ignore_index=True)

print(len(all_tiles_gmm12))
print(all_tiles_gmm12.shape)
```

```
106423555
(106423555, 1)
```

[11]:
```
all_tx['gmm12'] = all_tiles_gmm12['gmm12']
```

[16]:
```
# Define colors
k_colors = [
    "#2f4f4f", "#000000", "#008000", "#4b0082", "#ff0000", "#ffd700",
    '#b5bd61', "#00ffff", "#0000ff", "#ff69b4", "#1e90ff", "#ffdab9", "#ff00ff"
]
```

[ ]:
```
def get_major_cluster(values,bin_thresh=10):
    if len(values) < bin_thresh:
        return np.nan  # Return NaN for empty bins
```

```
    result = mode(values, keepdims=True)  # Ensure the result is always␣
↪array-like
    return result.mode[0] if result.mode.size > 0 else np.nan

# Parameters
bin_width = 5  # Adjust bin size
x_bins = int((data['x_location'].max() - data['x_location'].min()) / bin_width)
y_bins = int((data['y_location'].max() - data['y_location'].min()) / bin_width)
```

##### 5.2 Summarise cluster ids in hexbins and plot

We create hex bins with width 5, and cluster ids of transcripts in each bin are aggregated using a major voting method. We visualise the hex bins and color them with the aggregated cluster ids.

[34]:
```
data = all_tx
```

[35]:
```
data
```

[35]:
```
           Unnamed: 0    transcript_id     cell_id  overlaps_nucleus  \
0            11147569  281487862195756  ceoldomj-1                 1
1            11147570  281487862858835  fgclmdcj-1                 0
2            11147571  281487862587825  feompcff-1                 1
3            11147572  281487862133359  ffakaifg-1                 1
4            11147573  281487862319403  ffbjgjio-1                 1
...               ...              ...         ...               ...
106423550   141525001  282291020496974  oiakpbfj-1                 1
106423551   141525002  282291020496985  oiakpbfj-1                 1
106423552   141525003  282291020496904  oiakpbfj-1                 1
106423553   141525006  282291020496935  oiakpbfj-1                 1
106423554   141525007  282291020496914  oiakpbfj-1                 1

          feature_name  x_location   y_location  z_location    qv fov_name  \
0                A2ML1   7547.4375   162.781250   22.578125  40.0      A13
1                A2ML1   7725.9062   177.343750   19.203125  40.0      A13
2                 AAMP   7502.2500    90.859375   20.250000  40.0      A13
3                 AAMP   7529.1562    87.687500   21.812500  32.0      A13
4                 AAMP   7531.5780   165.437500   22.031250  33.0      A13
...                ...         ...          ...         ...   ...      ...
106423550        GSK3B  10271.9530  7821.203000   22.671875  38.5      K18
106423551        HSPA8  10272.6875  7822.468800   21.953125  37.0      K18
106423552        NOL4L  10271.8750  7822.078000   22.812500  34.5      K18
106423553       PIK3CA  10272.9060  7822.547000   22.390625  37.5      K18
106423554        PRPF3  10271.3125  7823.890600   22.531250  40.0      K18

           nucleus_distance  codeword_index codeword_category  is_gene  gmm12
0                    0.0000           12771   predesigned_gene     True      6
1                    0.8125           14663   predesigned_gene     True      4
```

```
2                         0.0000            2394  predesigned_gene     True      6
3                         0.0000            2394  predesigned_gene     True      6
4                         0.0000            2394  predesigned_gene     True     11
...                          ...             ...               ...      ...    ...
106423550                 0.0000            5051  predesigned_gene     True      6
106423551                 0.0000            9181  predesigned_gene     True      6
106423552                 0.0000            4147  predesigned_gene     True      6
106423553                 0.0000            9133  predesigned_gene     True      6
106423554                 0.0000            3271  predesigned_gene     True      6

[106423555 rows x 15 columns]
```

##### 5.3 The hex bin aggregated cluster visualisation

[36]:
```python
from matplotlib.colors import ListedColormap
```

[ ]:
```python
fig, ax = plt.subplots(figsize=(10, 8))

hb = ax.hexbin(
    data['x_location'],
    data['y_location'],
    C=data['gmm12'],  # Aggregation values
    gridsize=(x_bins, y_bins),
    reduce_C_function=get_major_cluster,  # Apply custom function
    linewidths=0.05,  # Thin line between bins
    cmap=ListedColormap(plt.cm.tab20.colors[:12]) # Set a discrete color palette
)

# Reverse y-axis
ax.invert_yaxis()

# Add labels and customizations
ax.set_xlabel("Ovarian Caner", fontsize=18)
ax.set_ylabel("Y Location", fontsize=18)
ax.tick_params(axis='both', which='both', colors='black')
ax.spines['top'].set_visible(False)
ax.spines['right'].set_visible(False)
ax.spines['left'].set_linewidth(1.5)
ax.spines['bottom'].set_linewidth(1.5)
ax.grid(False)

# Add color bar for hexbin
cb = fig.colorbar(hb, ax=ax, orientation='vertical')
cb.set_label("Major Cluster", fontsize=16)
```

[43]:
```python
fig.set_size_inches(14, 8)  # Set width and height in inches
fig
```

[43] :

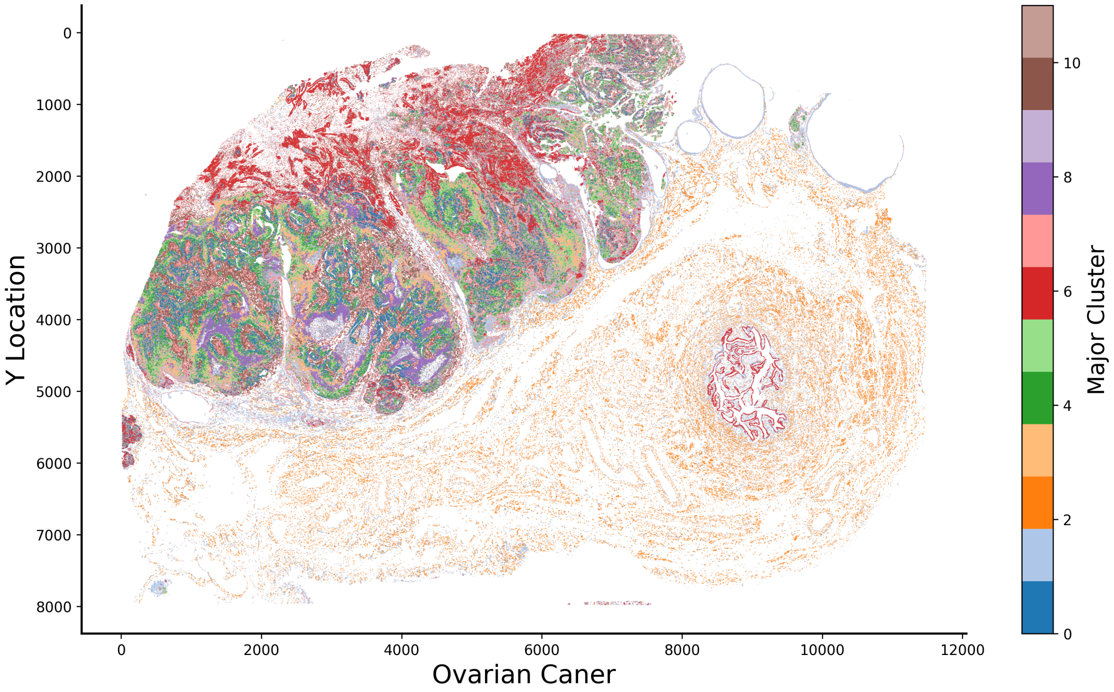
Y Location
Ovarian Caner
Major Cluster
0
1000
2000
3000
4000
5000
6000
7000
8000
0
2000
4000
6000
8000
10000
12000
0
2
4
6
8
10
